# Label-Free Electrochemical Imaging of Single Extracellular Vesicles

**DOI:** 10.1101/2024.05.21.595111

**Authors:** Yuanxin Zhang, Zichen Hong, Xuan Fu, Bo Yao, Guangzhong Ma

## Abstract

Extracellular vesicles (EVs) are pivotal in various biological processes and diseases, yet their small size and heterogeneity pose challenges for single EV quantification. We introduce interferometric electrochemical microscopy (iECM), a sensitive label-free technique that combines interferometric scattering and electrochemical impedance imaging. This method enables the quantification of the impedance of single EVs, providing unique insights into their electrochemical properties, and allows for the simultaneous measurement of size and real-time monitoring of antibody binding. Notably, we discovered that the impedance spectra of EVs can serve as unique fingerprints to differentiate EVs from cancerous and healthy cells, offering potential advancements in clinical diagnostics and the profiling of extracellular biomarkers.

## Introduction

Extracellular vesicles (EVs) are lipid bilayer-enclosed structures released by cells, garnering significant scientific interest due to their pivotal roles in diverse biological processes and disease pathways.^1,2^ The heterogeneous content of EVs, which includes proteins, nucleic acids, and metabolites, imparts distinct biological functions and potential diagnostic significance.^1^ However, accurately quantifying these vesicles presents substantial challenges due to their small size and complex heterogeneity.^3-5^

Fluorescence-based methods, such as high-resolution flow cytometry^6,7^ and fluorescence microscopy,^8-10^ have addressed the challenge of studying single EVs through multi-color immunostaining, enabling the profiling of various markers on individual EVs. Despite their advantages, these techniques are limited to detecting the presence of proteins rather than quantifying their abundance due to issues like labeling efficiency and photobleaching. In contrast, label-free techniques have emerged as valuable complementary tools for quantifying EVs.^11-13^ Nanoparticle tracking analysis (NTA) is widely used but lacks sensitivity and the ability to profile EVs.^14^ Techniques with high sensitivities, such as interferometric scattering (iSCAT),^15-17^ surface plasmon resonance (SPR),^18,19^ single particle interferometric reflectance imaging sensor (SP-IRIS),^20,21^ and dark-field microscopy,^22,23^ offer exceptional capabilities for imaging and sizing single EVs. However, they typically do not provide comprehensive profiles of single EVs either, as they measure only the binding of EVs to antibody-functionalized surfaces, thus limiting the detection to one type of protein per single EV. Surface-enhanced Raman spectroscopy (SERS) has been employed to probe the overall chemical content of single EVs,^24-26^ but it requires the use of non-uniform nanostructures for signal enhancement, which leads to signal inhomogeneity and reproducibility issues.^5^

Given the structural and compositional similarities between EVs and cells, electrochemical impedance spectroscopy (EIS) is an intuitive label-free biophysical approach for their analysis.^27^ Although impedance sensing and imaging methods have been utilized for measuring single cells^28-30^ and multiple EVs,^31,32^ achieving impedance imaging at the single vesicle level remains unprecedented due to both inadequate imaging sensitivity and the minimal electrochemical responses of single EVs. To address these challenges, we have developed a technique called interferometric electrochemical microscope (iECM). This method quantifies the electrochemical impedance of individual EVs while simultaneously measuring their size and membrane protein interactions. These key parameters allow us to distinguish EVs from different cell lines with high accuracy. Additionally, we demonstrate that by analyzing impedance spectra at specific fingerprint frequencies, EVs can be classified in a purely electrochemical manner. We believe our technique provides insights into the heterogeneity of EVs from a unique electrochemical perspective, contributing to our evolving understanding of these tiny vesicles.

## Results

### Setup and detection principle

The iECM setup, depicted in Figure 1a and S1, is built on an inverted microscope. The incident light is directed to the 60× oil immersion objective (NA = 1.50) and focused at the center of the back focal plane. The imaging substrate is an ITO-coated coverslip mounted with a silicone cell to secure the aqueous sample and electrodes in place. EVs attached to the surface are illuminated in wide-field mode, and the reflected light (from ITO) and the scattered light (from EV) are collected by a CMOS camera. An alternating potential is applied using a three-electrode configuration. Size measurements of EVs are performed using iSCAT imaging. This technique relies on the optical contrast generated by the interference between the scattered light and the reflected light. The image contrast, denoted as *C*_size_, is directly proportional to the cubic power of the particle diameter (*D*) based on the formula (see Methods)^15^

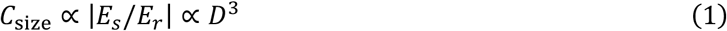

where *E*_r_ and *E*_s_ represent the fields of reflection and scattering, respectively. By measuring standard nanoparticle samples with known size to establish a calibration curve, the size of EVs can be determined from their *C*_size_.

**Figure 1.**
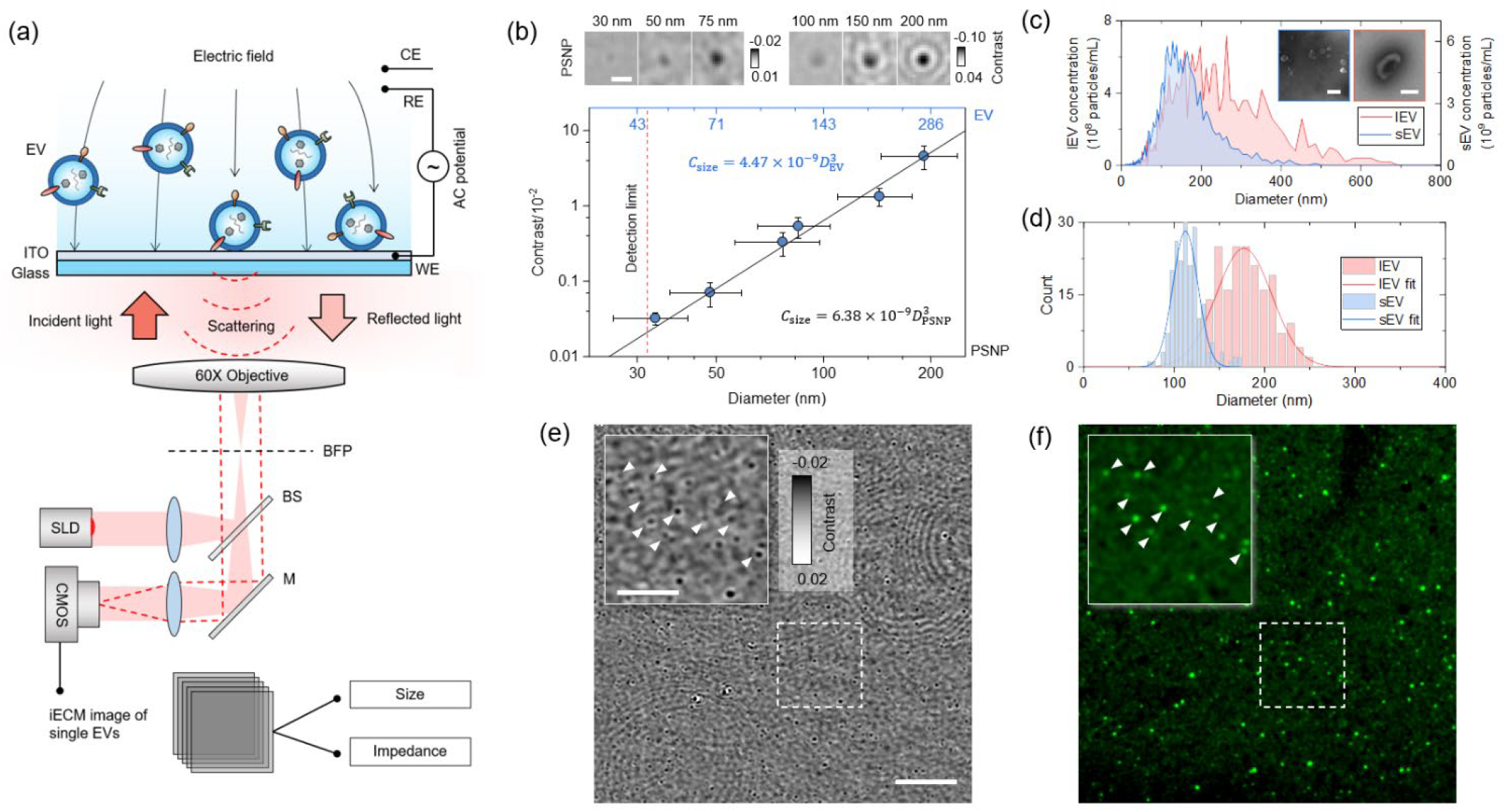
iECM imaging and sizing of EVs. (a) Schematic of the detection principle for simultaneous measurement of size and impedance of EVs. Size is determined by interferometric scattering (iSCAT) imaging, and impedance by electrical modulation applied to the surface. The electrochemical system consists of an ITO working electrode (WE), a Ag/AgCl quasi-reference electrode (RE), and a Pt coil counter electrode (CE). BFP, back focal plane; BS, 50/50 beam splitter; M, mirror. (b) Calibration curve for sizing EVs based on image contrast (*C*_size_) using polystyrene nanoparticles (PSNPs) with known diameters. The bottom x-axis indicates PSNP size, and the top x-axis shows the corresponding EV size producing the same image contrast. Inset: Example iSCAT images of PSNPs. Scale bar: 1 μm. (c) Size distribution of small EVs (sEVs) and large EVs (lEVs) characterized by nanoparticle tracking analysis (NTA). Insets show TEM images of sEV and lEV. Scale bars: 200 nm. (d) iSCAT-derived size distributions of EVs fitted to Gaussian curves. (e) iSCAT image of lEVs on the surface. Scale bar: 10 μm. Inset: Zoom-in of the marked square area. Scale bar: 5 μm. (f) Fluorescence image captured after iSCAT imaging at the same location, confirming the identity of spots as EVs. White arrows in insets of (e) and (f) indicate the same EVs. Rabbit CD63 antibody and Alexa Fluor 488-labeled goat anti-rabbit IgG were used for fluorescence imaging.

After measuring the size, an alternating voltage Δ*U* is applied to the ITO. This voltage modifies the surface charge density and, consequently, the dielectric constant of the ITO, leading to changes in ITO reflectance (Δ*Ref*_ITO_). In regions where EVs are present, the applied voltage on the ITO reduces from Δ*U* to Δ*U*′ due to the voltage drop caused by the EVs’ impedance. This reduction in voltage diminishes the local reflectance changes to Δ*Ref*_ITO+EV_, and the difference between Δ*Ref*_ITO_ and Δ*Ref*_ITO+EV_ creates image contrasts for the EVs. By modelling the solution/EV/ITO system with an equivalent circuit, the impedance density of the EVs (*Z*_EV_) can be quantified by (see Methods):

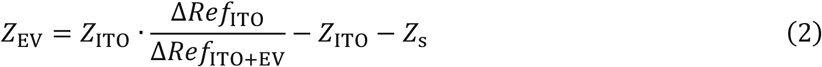

where *Z*_ITO_ and *Z*_s_ are the impedance density of the bare ITO and the solution, respectively. The ratio Δ*Ref*_ITO+EV_/Δ*Ref*_ITO_, denoted as impedance contrast (*C*_imp_), provides a direct measure of the EV’s impedance. We note that the scattered light from EVs is much weaker compared to the reflected light, thus its contribution to *C*_imp_ is negligible.

### Measuring the size of EVs

We first measured polystyrene nanoparticles (PSNPs) of various sizes using iSCAT to establish a size calibration curve. To discern the weak signal from the strong reflection, the first image, serving as the static background, was subtracted from subsequent images. Further noise reduction was achieved through moving average of the images, as detailed in the Methods section. The resulting image exhibited clear dark spots for PSNPs with diameter from 30 nm to 200 nm (Figures 1b and S2). The correlation between *C*_size_ and diameter was cubic, aligning with the theoretical predictions by Equation 1. Because PSNPs have higher refractive index than EVs, we converted the PSNP size into EV size to ensure accurate EV sizing (Figure 1b). Our setup achieved a detection limit of 32 nm for PSNPs and 45 nm for EVs, demonstrating sufficient sensitivity for most EV measurements (Supplementary Note 1).

EVs from A549 non-small cell lung cancer cells were separated into large extracellular vesicles (lEVs) and small extracellular vesicles (sEVs), which are originated from different biogenesis pathways.^33^ Their size, morphology, and protein expression were characterized by nanoparticle tracking analysis (NTA), transmission electron microscopy (TEM), and western blot respectively (Figures 1c and S3). Upon injecting 5 μL 10^8^/mL EVs into the silicone cell containing 495 μL PBS buffer, approximately 450 dark spots appeared within 6 seconds in a 120×120 μm^2^ region (Figure 1e and S4). To confirm the identity of these spots as EVs rather than cellular debris, we performed fluorescence imaging using the same setup (Figure S1). The EVs were incubated with antibodies targeting their membrane marker CD63, followed by labeling with Alexa Fluor 488-conjugated secondary antibodies. A comparison of the fluorescence and iSCAT images revealed excellent colocalization, confirming that the dark spots were indeed EVs (Figures 1e and 1f). Using the calibration curve, we sized hundreds of EVs from the iSCAT images, and the diameters of lEVs and sEVs were determined to be 177±44 nm and 112±20 nm, respectively. These measurements closely matched the NTA results, which recorded median sizes of 185 nm for lEVs and 132 nm for sEVs (Figures 1c and 1d).

### Electrical modulation of ITO

Next, we investigated the electrochemical imaging capability of iECM. We applied a sinusoidal potential (1 V at 20 Hz) to a bare ITO surface and recorded its mean reflectance at 500 frames per second (fps). The reflectance was modulated by the potential, exhibiting an oscillating profile (Figure 2a). To quantify the magnitude of Δ*Ref*_ITO_, we applied fast Fourier transform (FFT) to the oscillation curve over a one-second interval, which revealed a prominent peak at 20 Hz in the frequency domain (Figure 2b), corresponding to the amplitude of Δ*Ref*_ITO_ (see Methods). Using this approach, we extracted the Δ*Ref*_ITO_ for each pixel in the images, and generated a new image sequence (called iECM images) at a reduced frame rate of 1 fps, which contained spatially resolved Δ*Ref*_ITO_ data (Figure 2c).

**Figure 2.**
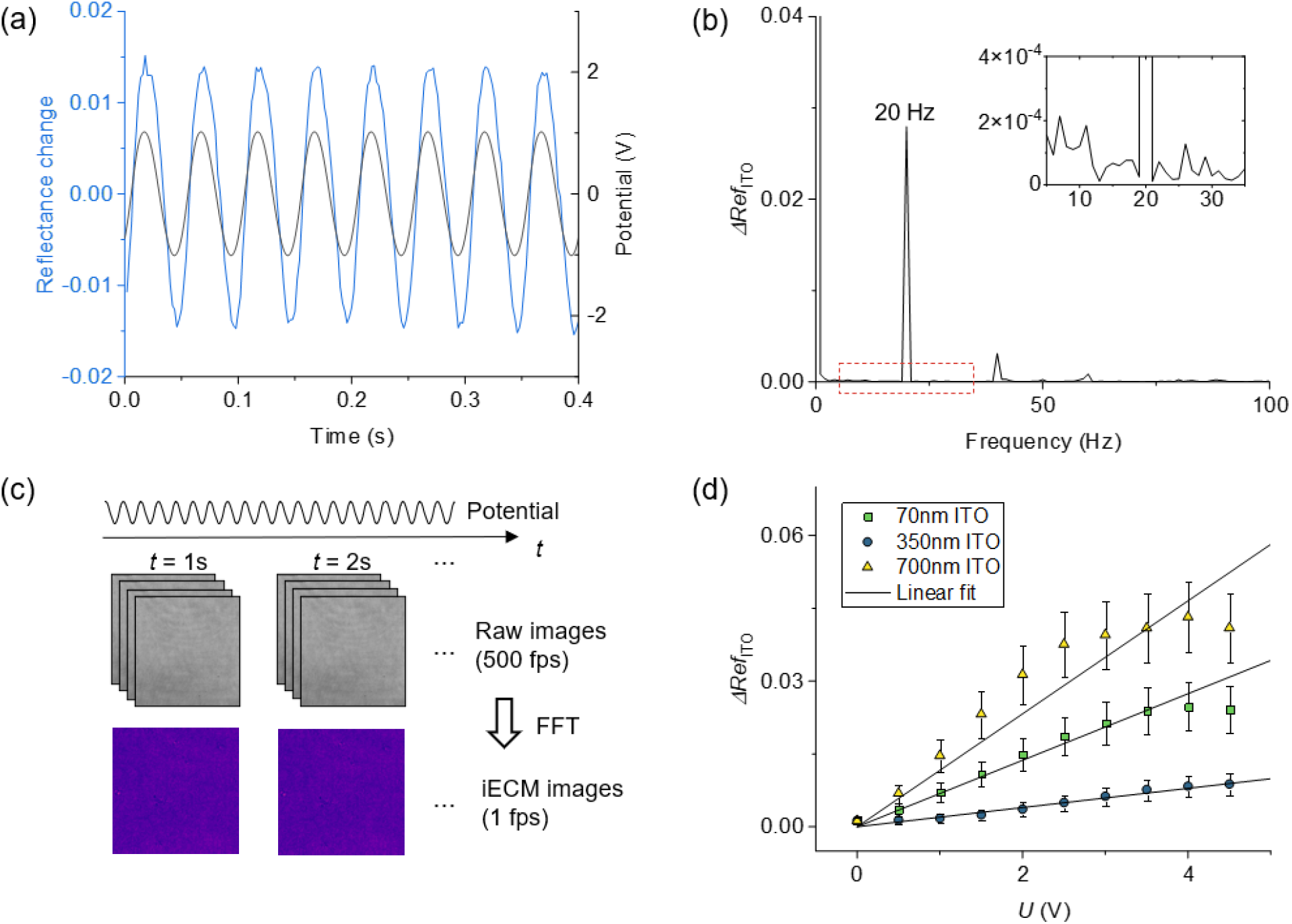
Electrical modulation of ITO. (a) Reflectance changes (Δ*Ref*_ITO_) of a 700 nm ITO film in response to an applied potential. The potential was a sinusoidal wave with a 1 V amplitude and 20 Hz frequency. (b) Fast Fourier transform (FFT) of Δ*Ref*_ITO_ over a 1-second period, with the peak at 20 Hz indicating the oscillation amplitude. The inset is an enlarge view of the squared region, showing minimal response at other frequencies. (c) iECM image processing. With the potentials applied, raw images are recorded at 500 frames per second. A 20 Hz FFT filter is applied to each pixel to generate an iECM image at a reduced frame rate of 1 fps. (d) Voltage response of ITO films of different thickness. Solid lines represent linear fittings of the data, and error bars show the standard deviation of Δ*Ref*_ITO_ measured from a 400×400 pixel area.

The sensitivity of Δ*Ref*_ITO_ to electrical modulation is critical for the performance of the sensing technique. We calculated the Δ*Ref*_ITO_ in Figure S5 and Supplementary Note 2 based on the Fresnel equation, and explored two important factors: the applied potential and the ITO film thickness. Our experimental result revealed a quasi-linear relationship between Δ*Ref*_ITO_ and applied potential, with distinct slopes for different ITO thicknesses (Figure 2d). These findings, corroborated by calculations, suggest that the 700 nm ITO film exhibits the highest sensitivity to potential changes. Consequently, this thickness was selected for impedance imaging in subsequent experiments.

### Impedance imaging of single EV

To image the impedance of EVs, we applied alternating potentials across a frequency range of 1 to 400 Hz to an ITO with attached EVs, and captured the corresponding iECM images. Most frequencies showed only background signals from the ITO, but notably, between 20-50 Hz, small spots emerged that resembled those in the iSCAT images (Figure 3a). These spots appeared only after injecting EVs onto the ITO (Figure S6) and closely colocalized with those observed in the iSCAT images (Figures 3b and 3c), confirming their identity as EVs rather than surface defects. We selected 31 single EVs and adjacent background regions to analyze their average Δ*Ref* across the frequency spectra (Figure 3d). Both regions exhibited decreasing Δ*Ref* with increasing frequency, but with different patterns. By calculating the ratio Δ*Ref*_ITO+EV_/Δ*Ref*_ITO_ from the curves, we observed a discernible peak in the low-frequency domain, where optimal iECM imaging contrast is achieved. Similar impedance patterns were previously observed in adherent cells,^34^ yet the optimal sensing frequency was located at 1 kHz, with a broader peak ranging from 100 Hz to 100 kHz. The distinct response in EVs can be attributed to their unique electrochemical properties compared to whole cells (Supplementary Note 3). It is noteworthy that while most EVs in the iSCAT image corresponded to those in the iECM images, discrepancies were observed due to a few additional free EVs binding to the surface after iSCAT video recording. Also, some EVs failed to establish sufficient contact with the surface, thereby not contributing to the impedance within the electrical circuit.

**Figure 3.**
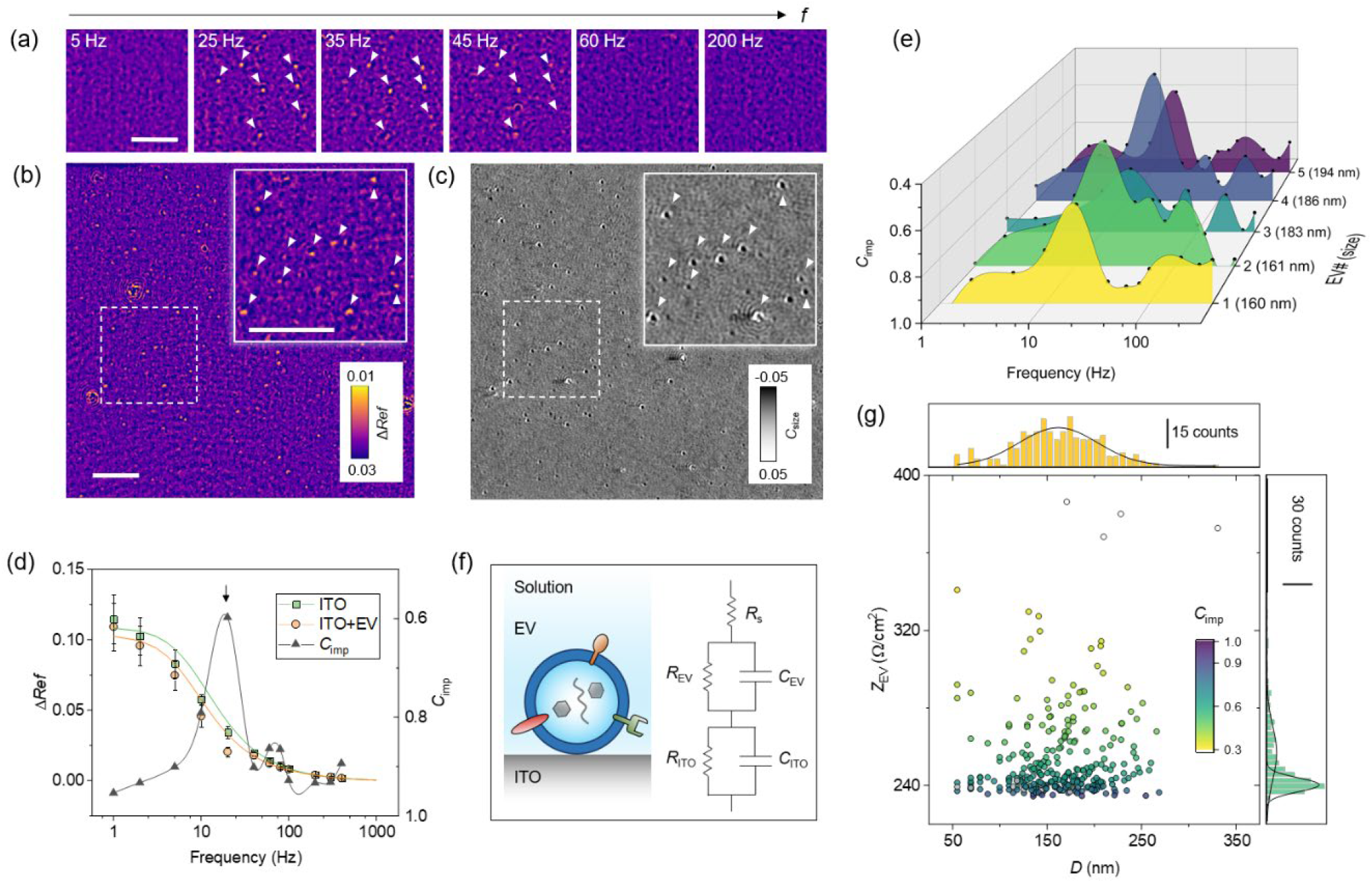
Impedance imaging of single EVs. (a) Example iECM images showing the frequency response of EVs. Contrast has been optimized for clarity. Several individual EVs are indicated with white arrows. Scale bar, 5 μm. (b) iECM image at 20 Hz, displaying a 75 μm×75 μm region with approximately 100 single EVs. Inset: Closer view of the square-dashed region. Scale bars: full image, 10 μm; inset, 5 μm. (c) iSCAT image of the same region as in (b). White arrows in the insets of (b) and (c) mark the same EVs. (d) Reflectance change (Δ*Ref*) across different frequencies for bare ITO region and EV region (ITO+EV). Error bars represent standard deviations for 31 background regions and 31 individual EVs, respectively. Solid curves are simulation results using the model described in (f). The black curve shows the ratio Δ*Ref*_ITO+EV_/Δ*Ref*_ITO_ with the peak for optimal iECM image contrast marked by an arrow. Details about the simulation are described in Supplementary Note 4. (e) Impedance spectrum for five individual EVs. (f) Equivalent circuit model for EV impedance imaging. *R*_s_ is the solution resistance; *R*_EV_ and *C*_EV_ are the EV resistance and capacitance, respectively; *R*_ITO_ and *C*_ITO_ are the ITO interfacial resistance and capacitance, respectively. (g) Size and impedance of 295 single EVs, color-coded by *C*_imp_ value. Data are projected onto the x- and y-axes, showing size and impedance distributions, which are fitted with Gaussian curves.

Subsequent analysis focused on the response from individual EVs. Figure 3e shows five single EVs of varying sizes, displaying differences in their frequency spectra with peaks between 20 Hz and 50 Hz, which reflects their electrochemical heterogeneity. To figure out the electrical contribution of EVs in the measurement circuit, we adapted a model based on Randle’s equivalent circuit, which contains contributions from the solution, the EV, and the ITO electrode (Figure 3f). Given the structural similarity of EVs to cells, we modeled each EV with a resistor and a capacitor in parallel, representing the lumen and the membrane, respectively. Using this model, we simulated the frequency spectra for ITO and ITO+EV, which closely matched the observed data (Figure 3d, solid curves). More importantly, this model allows us to extract the impedance for individual EVs. Using Equation 2 and measured resistance and capacitance in the circuit (Figure S7), *Z*_EV_ can be directly derived from *C*_imp_ in the iECM image. Note that we used impedance density throughout the calculation, therefore the unit for *Z*_EV_ is Ω/cm^2^. We determined the impedance density (at *f* = 30 Hz) of 295 individual EVs, and plotted them against their sizes (measured by iSCAT) on a map (Figure 3g). No clear correlation or clusters were evident (Figure S8), suggesting that impedance density is independent from the size. Unlike the single size peak observed at 163 nm, the impedance showed two peaks at 240 Ω/cm^2^ and 257 Ω/cm^2^, indicating two distinct electrical states within the EV population. These variations may be attributed to differences in structural factors such as conformation, density, and surface charge, as well as compositional diversity in proteins, nucleic acids, metabolites, and lipids.^3^ Further elucidating the roles of these factors calls for the integration of iECM with other advanced analytical techniques, which is beyond the scope of this work. However, our use of direct stochastic optical reconstruction microscopy (dSTORM) has revealed significant heterogeneity in surface protein content. Using antibodies against EpCAM, CD63 and CD81, which are three EV membrane proteins, dSTORM revealed single positive, double positive, or triple positive expressions of the proteins in single EVs (Figure S9).

### Antibody binding and EV profiling

Antibody binding to the EV membrane introduces impedance changes at the interface, which are detectable by iECM in real-time (Figure 4a). We quantified the interactions between A549-derived lEVs and EpCAM, CD63, and CD81 antibodies. We initially established an impedance baseline in PBS buffer over 60 seconds. Then 10 nM EpCAM antibody was introduced into the system, followed by a 2-minute incubation period. This procedure was repeated for 6.6 nM CD63 and 6.6 nM CD81 antibodies. Figures 4b and 4c illustrates the impedance changes for three representative single EVs (EV#6-8), with a 200 nm PSNP and a blank ITO region as negative controls. The EVs displayed impedance responses, while PSNP and bare ITO showed no change. Notably, the response varied among EVs: EpCAM antibody induced a negative impedance change in EV#6 but no change in EV#7 and EV#8, while CD63 and CD81 antibodies caused positive changes in EV#7 and EV#8. This difference is attributed to the varying expression levels of the markers. The observation was confirmed by measuring multiple EVs simultaneously (Figure S10). Figure 4b also shows that the impedance fluctuations for EVs were higher than those for PSNPs and the background (Figure 4d), likely reflecting the soft and flexible nature of the EVs.^35^

**Figure 4.**
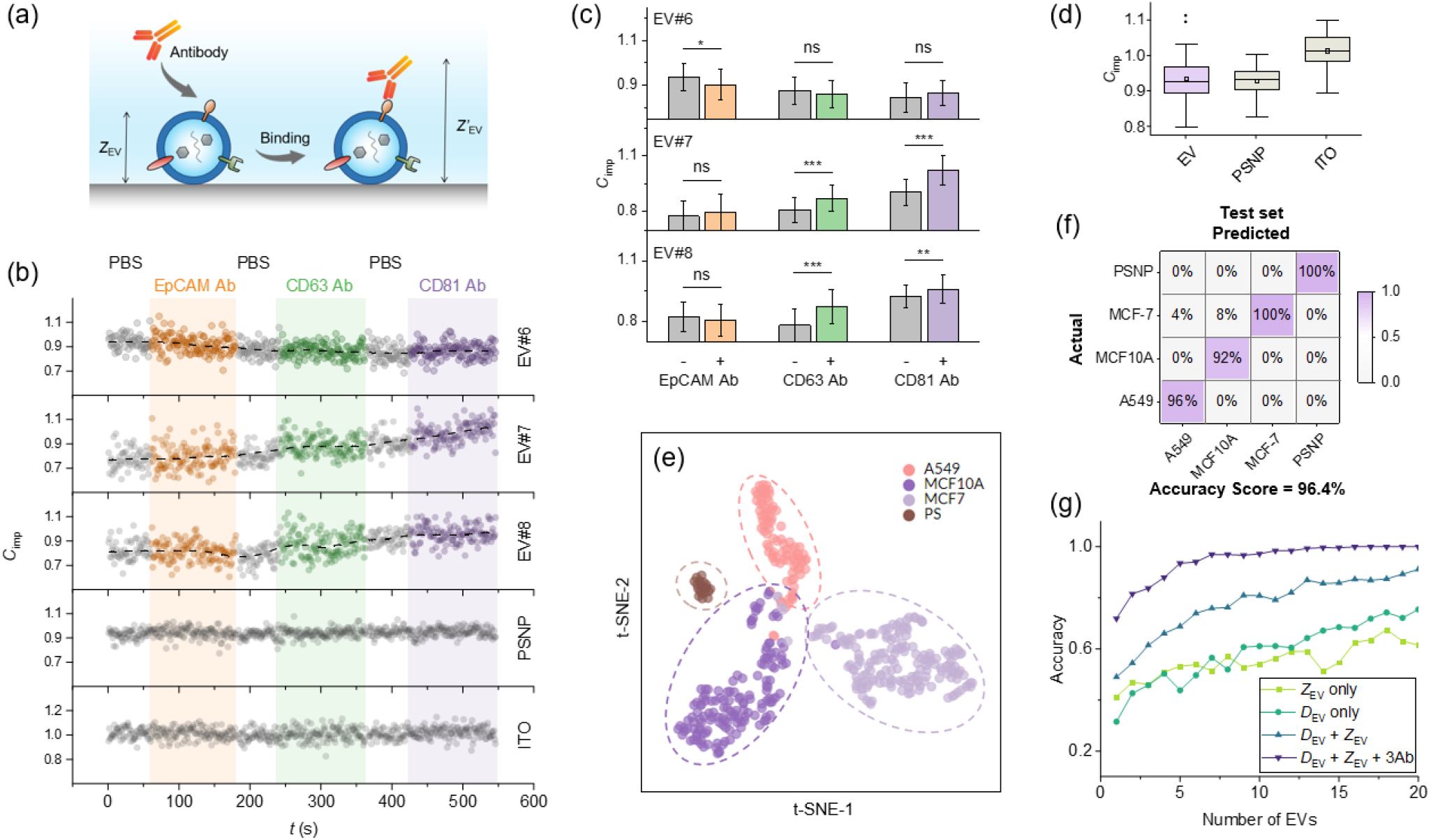
Label-free EV profiling. (a) Schematic illustration of antibody binding to EV membranes. (b) Real-time impedance monitoring of three A549 EVs upon antibody interaction. Antibodies specific to EpCAM, CD63, and CD81 were sequentially introduced, with each binding event lasting 2 minutes, as depicted by the shaded areas. A representative PSNP and a background ITO region served as controls. The applied frequency was set at 30 Hz. (c) Statistical analysis of impedance changes before and after each antibody binding. Impedance contrasts measured 1 minute before and 1 minute after introducing each antibody (60 data points each) were compared. Statistical significance is denoted as *:p<0.05, **:p<0.01; ***p<0.001; ns = not significant. (d) Temporal fluctuation of impedance. Impedance contrasts values (continuously recorded for 1 minute) for a representative A549 EV, a PSNP, and a background ITO region are presented in a box plot. (e) t-SNE plot of EVs from different cell lines and PSNPs. The analysis is based on size, impedance (at 30 Hz), and antibody-induced impedance changes (by EpCAM, CD63, and CD81 antibodies, respectively). Each data point represents the averaged value from a randomly selected set of 10 EVs, a method employed for noise reduction. (f) The confusion matrix of the test set when using the Random Forest algorithm to classify EVs. Data were collected on three different ITO chips for each type of EV. (g) Relationship between classification accuracies (training set) and the number of EVs used for averaging. Generally, increased averaging enhances accuracy. The curves illustrate accuracies derived from either individual parameters or a set of parameters. *D*_EV_, diameter; *Z*_EV_, impedance; 3Ab, with EpCAM, CD63, and CD81 antibodies.

The capability of simultaneously measuring impedance, antibody binding and size for the same single EV allows for identifying different types of EVs. To demonstrate this, we studied the EVs derived from A549 cells, MCF-7 breast cancer cells, and MCF10A normal breast epithelial cells, by measuring their size (*C*_size_), impedance at 30 Hz (*C*_imp_), and antibody binding induced *C*_imp_ changes (including EpCAM, CD63 and CD81 antibodies) (Figure S11). These 5-dimensional data were plotted in a 2D t-SNE plot (Figure S12). Initial trials using raw data from single EVs failed to achieve clear classification, likely due to high temporal fluctuations in *C*_imp_ and biological similarities in EV structure. To reduce the noise, we employed a bootstrap averaging method: for each EV, we randomly selected another nine EVs of the same type and computed the mean impedance for these ten EVs. These mean values were then used for t-SNE classification, which successfully produced distinct clusters corresponding to EVs from different sources (Figure 4e). The same procedures were applied to PSNPs as controls, which yielded a separate cluster. We further developed machine learning models using Random Forest (RF) and Deep Neural Networks (DNN) for classification, which could achieve overall accuracy up to 99.4% of train set and 100% of test set for RF (Figure 4f and Supplementary Figure S11) and both 100% for DNN (Supplementary Figure S13) when distinguishing groups of 10 EVs. The accuracy improves with the inclusion of more EVs in the bootstrap group, which escalates to near 100% when data from about seven EVs are pooled (Figure 4g). We also examined the classification accuracies using only *C*_size_, *C*_imp_, or a combination of both, but they resulted in lower accuracies compared to the combined data approach. Together, these analyses suggest that aggregating data from multiple EVs is necessary for reliable classification.

### Electrochemical profiling of EVs

The impedance spectrum of EVs in Figure 3e displays unique patterns within the 20-50 Hz range, suggesting the possibility to use this fingerprinting region alone for EV identification without relying on size or antibodies. To investigate this, we measured the impedance spectra at five frequencies (45, 40, 35, 30, 25 Hz) for different EVs and PSNPs, and presented the results in Figure 5a. Each data point represents the average *C*_imp_ of multiple single EVs or nanoparticles. The distinct spectra among the samples indicate intrinsic electrochemical differences. The normalized impedance for individual EVs and PSNPs is illustrated in a heatmap in Figure 5b. Using the bootstrap averaging method, we mapped the *C*_imp_ values from groups of 10 EVs onto a t-SNE plot, which resulted in clear separations among the samples (Figure 5c). Background regions on the ITO without EVs overlapped on the t-SNE plot, suggesting that the separation of EVs is not due to measurement errors between different samples. Both RF and DNN machine learning models yields 80-90% accuracy (Figures 5d, S13, and S14). These findings underscore the capability of iECM in distinguishing different EV populations using as few as 10 EVs, based solely on their electrochemical properties.

**Figure 5.**
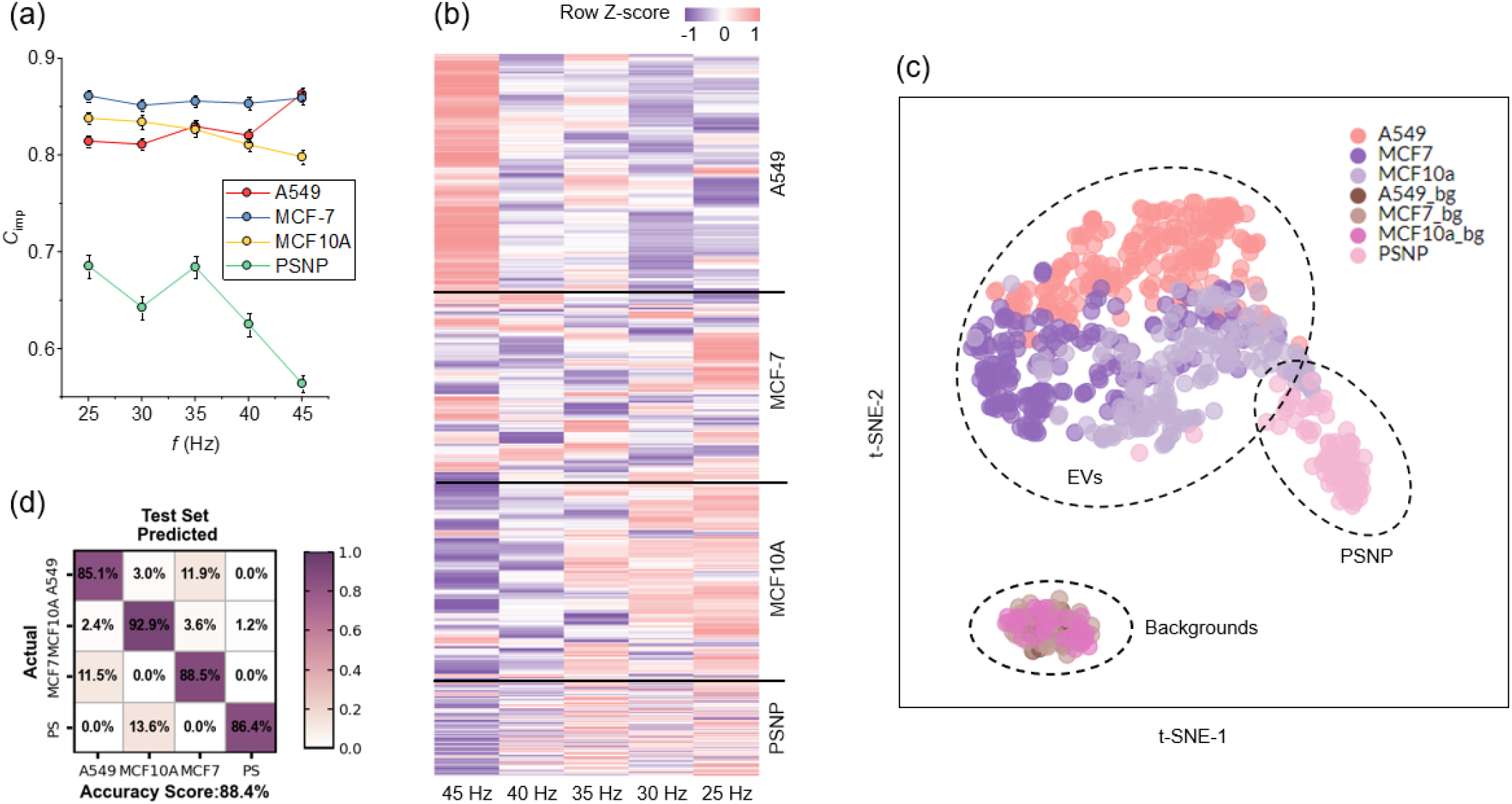
Electrochemical impedance profiling of EVs. (a) Impedance spectra of EVs from different cell lines (A549, MCF-7, MCF10A) and PSNPs within the fingerprint frequency domain from 25 Hz to 45 Hz. The data points represent average impedance contrast from 204 A549 EVs, 178 MCF-7 EVs, 171 MCF10A EVs, and 88 PSNPs; error bars indicate standard error. (b) Heatmap displaying the impedance contrast values for each individual EV or PSNP shown in (a). Data are normalized row-wise for enhanced visualization. (c) t-SNE mapping of EVs, PSNPs, and background regions randomly selected on the iECM images (30 background regions for each cell type, labeled by _bg). Each EV data point represents the average value from a randomly selected set of 10 EVs. (d) The confusion matrix of the test set when using the Random Forest algorithm to classify EVs. Data were collected on three different ITO chips for each type of EV.

## Discussion

Our study presents the iECM as a useful addition to the existing arsenal of label-free techniques for analyzing EVs. Through the development and application of iECM, we have demonstrated its ability to quantify the impedance and size of individual EVs and to assess molecular interactions at the single-vesicle level. This method distinguishes EVs from various cell types based on their unique electrochemical signatures, offering a novel approach for EV classification and providing valuable insights into their biophysical properties.

Most existing EV profiling methods, such as fluorescence and SERS, primarily focus on the biochemical properties of EVs. In contrast, iECM exploits electrical (and size) signatures, offering a distinct biophysical approach to EV profiling that complements these existing methods. Similar to how the Raman spectrum leverages vibrational energy features, iECM utilizes the impedance spectrum to convey specific electrical responses inherent to the EV structure. This approach has potential benefits, particularly if the resolution and frequency range of the technology can be further improved. Moreover, when combined with the latest advancements in iSCAT, which can now measure single biomolecules down to the 10 kDa level,^36^ iECM could extend its applications to probe single electro-active molecules, such as ion channels, lipoproteins, electron transfer proteins, and DNA.^37^ These technological enhancements could significantly advance the fields of EV and single-molecule research, providing more detailed and precise characterizations of these crucial biological entities.

## Methods

### Experimental setup

The setup was built on an inverted microscope (Olympus IX-73) equipped with a 60× (NA 1.50) oil-immersion objective. A superluminescent light-emitting diode (SLED) with a central wavelength of 670 nm (SLD-260-HP, Superlum) was used as the light source. The output power utilized for iSCAT and impedance imaging was 10 mW, controlled by a SLED current driver (PILOT4-AC, Superlum). The incident light was collimated and then focused onto the back focal plane of the objective to facilitate wide-field illumination. A high-speed CMOS camera (mini2MGE-CM4, Mikrotron) was used to record images. For electrical modulation, a three-electrode configuration was used, comprising an indium tin oxide (ITO) as the working electrode, an Ag/AgCl wire as the quasi-reference electrode, and a Pt coil as the counter electrode. The potential was applied via a potentiostat (WaveDriver 200, Pine) in conjunction with a function generator (DG1022Z, Rigol). Synchronization between the camera and the potentiostat was achieved through a data acquisition card (USB-6003, NI).

### Materials

ITO coated coverslips with different thickness (70, 350, and 700 nm) were purchased from SPI Supplies. According to the manufacturer, these coverslips have a refractive index of 2 and an extinction coefficient of 0.0044 at 650 nm. Polystyrene nanoparticles (PSNPs) with diameters of 30, 50, 75, 100, 150, and 200 nm were acquired from Feynman Nano. Their actual sizes were confirmed using dynamic light scattering (Zetasizer Nano ZS90). Alexa Fluor 647 anti-human CD63 antibody (353016) was obtained from BioLegend. Alexa Fluor 488-labeled mouse anti-human CD81 antibody (MCA1847A488) was purchased from Bio-Rad. Alexa Fluor 555-labeled mouse anti-human EpCAM antibody (5488S) was from Cell Signaling Technology. Rabbit CD63 antibody (25682-1-AP) was purchased from Proteintech. The corresponding secondary antibody, Alexa Fluor 488-labeled goat anti-rabbit IgG (A0423), was obtained from Beyotime. RIPA lysis buffer was from Sangon Biotech. Penicillin-streptomycin solution (100×P/S), phosphate-buffered saline (1×PBS), and trypsin-ethylenediamine tetraacetic acid (Trypsin-EDTA) solution were obtained from Beyotime. MCF-10A, MCF-7, A549 cell lines, and specific culture medium for MCF-10A and MCF-7 were purchased from Pricella. RPMI 1640 medium was from VivaCell.

### EV preparation and characterization

A549 cells were cultured in 1640 medium supplemented with 10% fetal bovine serum (FBS) and 1% P/S. MCF-10A cells were maintained in specific culture medium consisting of DMEM/F12, 5% horse serum, 20 ng/mL epidermal growth factor (EGF), 0.5 μg/mL hydrocortisone, 10 μg/mL insulin, 1% non-essential amino acids (NEAA), and 1% P/S. MCF-7 cells were cultured in MEM (with NEAA) supplemented with 10 μg/mL insulin, 10% FBS, and 1% P/S. All cell lines were incubated at 37°C with 5% CO_2_. Upon reaching 70%-80% confluency, the supernatant was removed and the cells were rinsed with 1×PBS. Subsequently, the cells were cultured for 48 hours in their respective serum-free media, and the supernatant (supernatant A) was then collected for EV isolation. The supernatant A was centrifuged at 2000×g for 10 min to remove dead cells and cell debris, followed by 10000×g for 0.5 h to obtain precipitate A and supernatant B. The precipitate A, namely large extracellular vesicle (lEVs), was resuspended in 1×PBS and stored at -80 °C for future use. The supernatant B was filtered with a 220 nm filter (Millipore) and centrifuged at 150000×g for 2 h (Beckman XPN-100 Coulter) to obtain precipitate B, namely small extracellular vesicle (sEVs). Then the sEVs were washed with 1×PBS and centrifuged again at 150000×g for 2 h for purification. Finally, the sEVs were resuspended in 1×PBS and stored at -80 °C for future use. The size of the EVs was characterized using nanoparticle tracking analysis (NTA) and transmission electron microscopy (TEM). The expression of membrane proteins was confirmed with western blot (Figure S3) and super-resolution microscopy (Figure S9). Both sEVs and lEVs were used for iSCAT sizing measurement, while only lEVs were used for impedance related measurements.

### Size imaging

The ITO surface was cleaned with ethanol and water, each for three times, and dried with nitrogen. For nanoparticle imaging, the nanoparticle stock solution was diluted 100 times. For EV imaging, a final concentration of 10^8^ EV particles/mL in PBS was used. Videos capturing the interaction of PSNPs or EVs with the surface were recorded, with camera settings detailed in Table S1. Initially, temporal noise reduction was applied by averaging 11 consecutive frames for particles larger than 75 nm, and 51 frames for particles smaller than 50 nm and for EVs. Each frame was then spatially smoothed using Fiji’s smooth function. The first image (N_0_) of the series (N_0_, N_1_, …, N_n_) was subtracted from each subsequent frame as static background. The differential sequence was divided by N_0_ to create a contrast image sequence, S_n_ = (N_n_ – N_0_)/N_0_, where nanoparticles appear as dark spots. In iSCAT, the light intensity *I* recorded by the camera includes the contributions from both reflected and scattered light (*E*_*r*_ and *E*_*s*_), described by

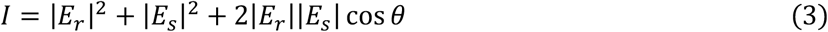

where *θ* is the phase difference between the two fields. The static background of iSCAT image, *I*_*bg*_, is dominated by |*E*_*r*_ |^2^. Given the weak scattering term from small nanoparticles, |*E*_*s*_|^2^ is considered negligible. Consequently, the image contrast (*C*_size_) is defined as:

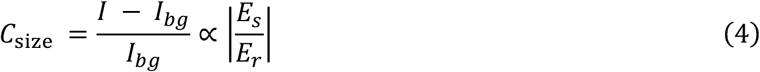

For our experiment, a diffraction limit-sized region (3×3 pixels) was used to measure the contrast for each nanoparticle or EV. The ratio |*E*_*s*_/*E*_*r*_| is proportional to the polarizability α (and thus *D*^3^) of the particle (Supplementary Note 1), leading to *C*_size_ ∝ *D*^3^.

The detection limit for sizing is constrained by the noise level, which is quantified by the standard deviation of intensity within a diffraction-limited region after spatial and temporal smoothing. Currently, our setup achieves a detection limit of 45 nm for EVs, which adequately covers the size range of EVs.

### Impedance imaging

Electrochemical measurements were performed immediately after the EVs settled on the surface. Unbound EVs were removed by flushing the surface with 5 times diluted PBS solution, which also served as the medium for subsequent impedance imaging experiments. Next, a sine wave was applied to the ITO, which changes the dielectric constant of the ITO (*ε*_ITO_) and hence the refractive index 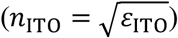. The relationship between the dielectric constant change and the applied voltage is described by:

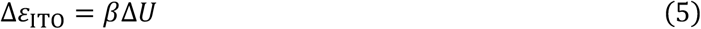

where *β* is a constant dependent on the physical properties of the ITO film, given by 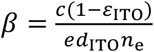 (Supplementary Note 2). Here, *R, n*_e_, *c*, and *d*_*ITO*_ represent the elementary charge, charge density, interfacial capacitance density, and the thickness of the ITO film, respectively. As a result, the ITO reflectance *Ref*_ITO_ varies with the voltage-modulated dielectric constant, following the equation:

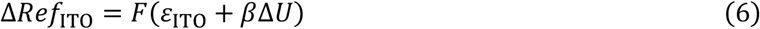

The function *F* corresponds to the Fresnel equation for the three-layer system of glass/ITO/water. In the ITO region where EVs are present, the applied voltage Δ*U* diminishes to Δ*U*′ due to the voltage drop across the EVs, influencing local reflectance as:

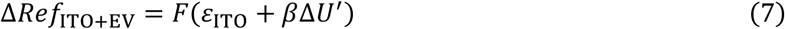

Using the quasi-linear relationship between Δ*Ref*_ITO_ and Δ*U* established in Figure 2d, we have Δ*Ref*_ITO_ = *k*Δ*U* and Δ*Ref*_ITO+EV_ = *k*Δ*U*′. These relationships simplify the results of the Fresnel equation, enabling the calculation of the impedance contrast *C*_imp_:

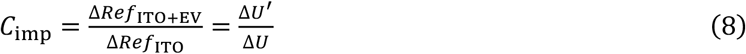

By employing the equivalent circuit model in Figure 3f, we can deduce the relationship between *U* and *U*^′^:

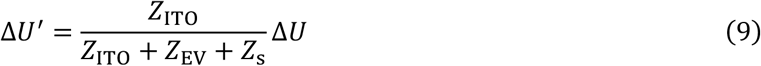

where *Z*_ITO_, *Z*_EV_, and *Z*_s_ are the impedance for bare ITO, EV, and solution, respectively. Combining Equations 8 and 9 yields Equation 2 in the main text. We note that although *C*_imp_ does not depend on the absolute amplitude of Δ*U*, Δ*U* must be sufficient to induce a Δ*Ref*_ITO_ detectable above optical noise levels. It is important to carefully control the applied voltage to optimize the signal-to-noise ratio without damaging the ITO (Table S2).

With the alternating potential applied to the ITO, the images were recorded at a frame rate of 500 fps with an exposure time of 1.5 ms. A fast Fourier transform (FFT) filter processed the recorded images over one second to extract signal amplitude (*I*_FFT_) and phase at the set frequency. The reflectance change for each pixel was calculated by Δ*Ref* = *I*_FFT_/*I*_0_, where *I*_0_ represents the initial pixel intensity without applied potential. This method produced iECM images such as Figure 3b, which displays the spatially resolved Δ*Ref*. To calculate the Δ*Ref*_ITO+EV_ of the EVs in the image, we applied a Gaussian blur (sigma = 3) to minimize the visibility of small EV spots while preserving the ITO background. The original image was then divided by the blurred image (Equation 8), whereby each pixel in the resulting image represents the *C*_imp_. The *C*_imp_ value for each EV can be directly measured from this image.

The relationship between *C*_imp_ and EV’s impedance density is shown in Figure S16. The *C*_imp_ values from different ITO chips were analyzed to ensure that measurement accuracy remained consistent (Figure S17). The measured impedance signal can be influenced by the non-uniform electro-optical response of the ITO, including both spatial and temporal noises (Figure S18). We found that the overall spatiotemporal noise is 0.137 in terms of *C*_imp_, corresponding to 0.31 Ω/cm^2^, which determines the impedance detection limit.

## Acknowledgments

We acknowledge financial support from the National Natural Science Foundation of China under grant numbers 22374127 and 22174126, and the startup funding from Zhejiang University. We thank Cheng Ma from the Core Facilities, School of Medicine at Zhejiang University for help in ultracentrifugation; and supports from the Chemistry Instrumentation Center at Zhejiang University in dynamic light scattering and TEM measurements.

## Author contributions

G. M. and B. Y. conceived and supervised the project. G. M. designed the experiments and implemented the instruments. Y. Z., Z. H., and G. M. carried out the iECM experiments and analyzed the data. Z. H. and X. F. contributed in EV preparation and characterization. G. M., B. Y, and Z. H. wrote the paper.

## Additional Information

### Competing interests

The authors declare no conflict of interest.

**Supplementary Information** is available for this paper. Supplementary Figures 1–18, Tables 1–2 and Notes 1–5.

### Code availability

The code and data of this study are available from: https://github.com/HgZnCH3/EV_impedance.

**Correspondence and requests for materials** should be addressed to G.M.

