## Supplementary Information for "Label-Free Electrochemical Imaging of Single Extracellular Vesicles"

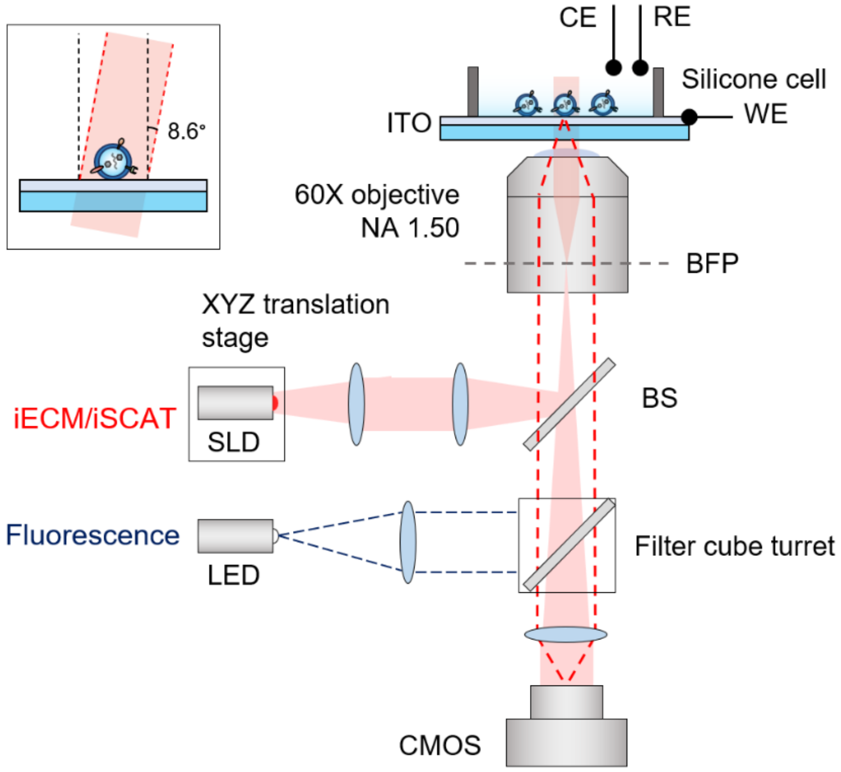


**Supplementary Figure 1. iECM setup.** The setup is based on an Olympus IX73 two-deck system, where the upper deck is used for the iECM/iSCAT channel and the lower deck for the fluorescence channel. For iECM/iSCAT imaging, the filter cube turret on the lower deck is switched to an empty position, allowing transmission of the reflected and scattered light (dashed line) to the camera. The incident light is tilted approximately 8.6° to minimize glares by slightly moving the focus position away from the center of the back focal plane (BFP) using the translation stage. This small tilting angle does not affect the sensitivity of electrochemical imaging (see Figure S5a for details).

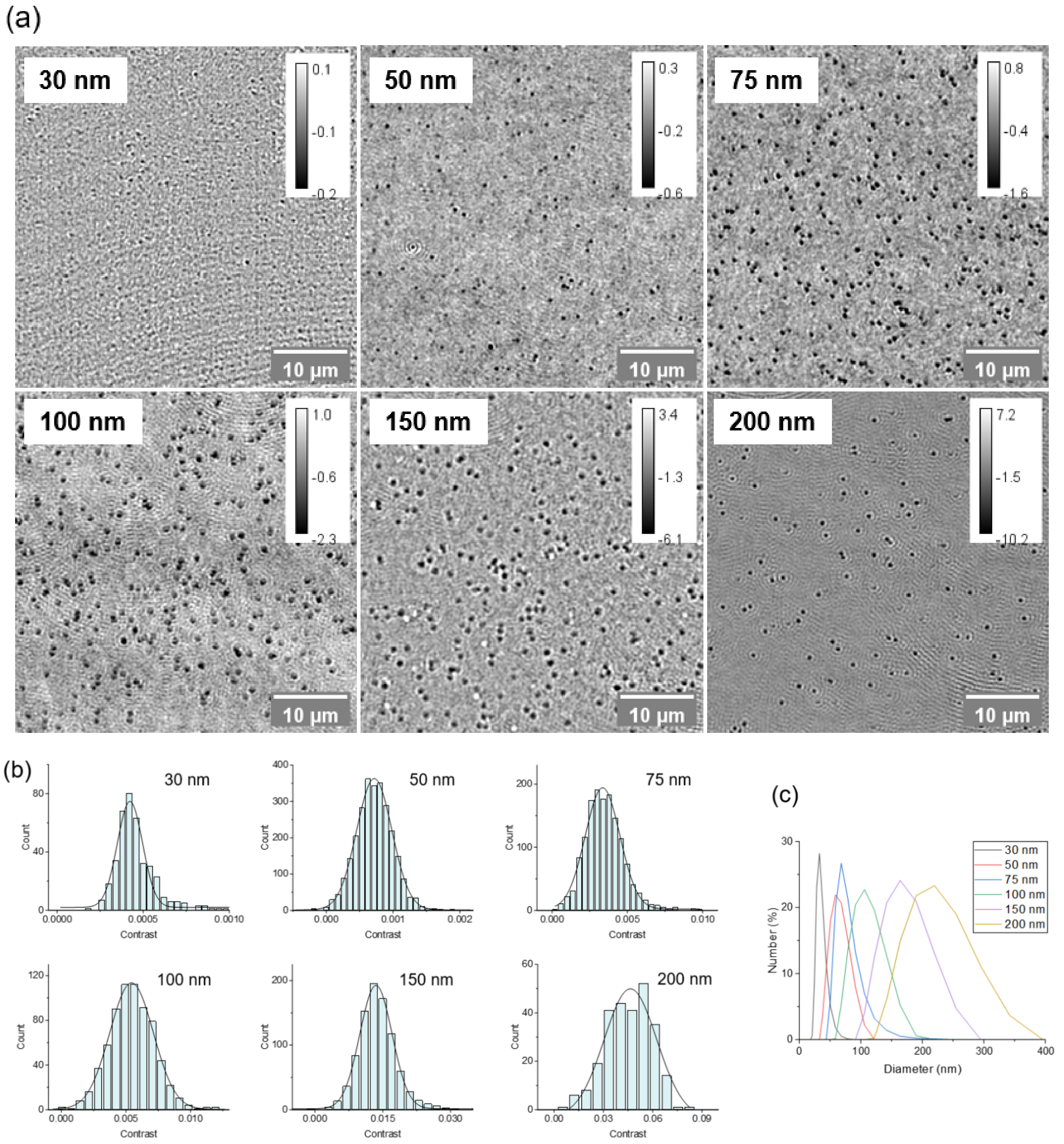


**Supplementary Figure 2. Establishing size calibration curve using polystyrene nanoparticles.** (a) iSCAT images of nanoparticles of different sizes. Calibration bars indicate grayscale levels. Experimental conditions can be found in Table S1. (b) Contrast distribution for the nanoparticles. The solid curves represent Gaussian distribution fittings. (c) The actual sizes of the PSNPs were measured using dynamic light scattering.

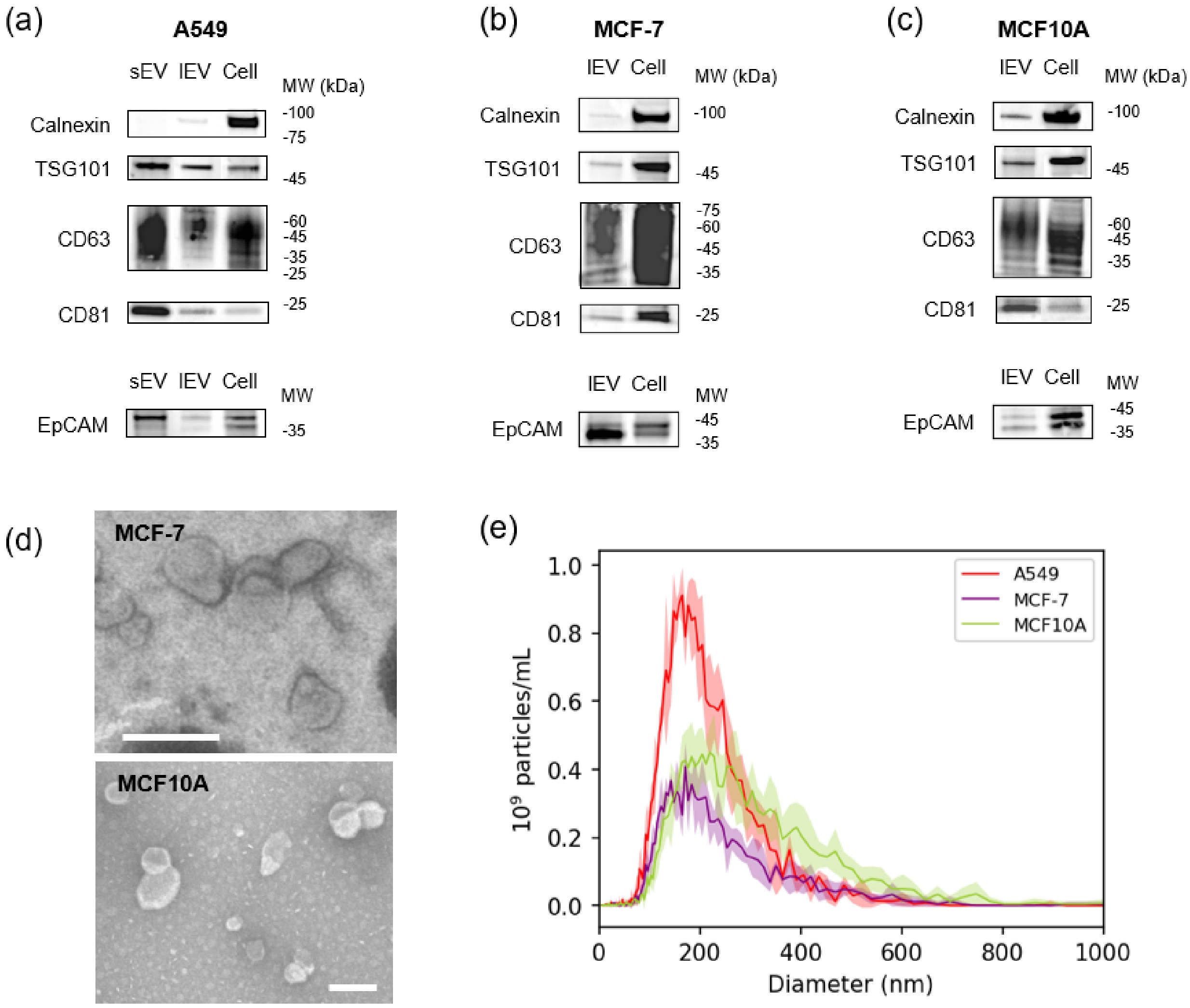


**Supplementary Figure 3. Characterization of EVs.** (a-c) Western blot analysis for A549, MCF-7, and MCF10A cells and their sEVs and/or lEVs. Three positive protein markers (TSG101, CD81 and CD63) and one negative protein marker (Calnexin) were selected for sEVs as recommended by the MISEV2018 guidelines.[1] The presence of positive protein markers and the absence of Calnexin in (a) confirm the successful isolation of sEVs. The presence of EpCAM in both sEVs and lEVs is also verified. (d) TEM images of lEVs from MCF-7 and MCF10A cells. Scale bars: 250 nm. (e) Size distribution of lEVs measured by NTA. The solid curves represent mean values with shaded region indicating standard deviation from at least four measurements.

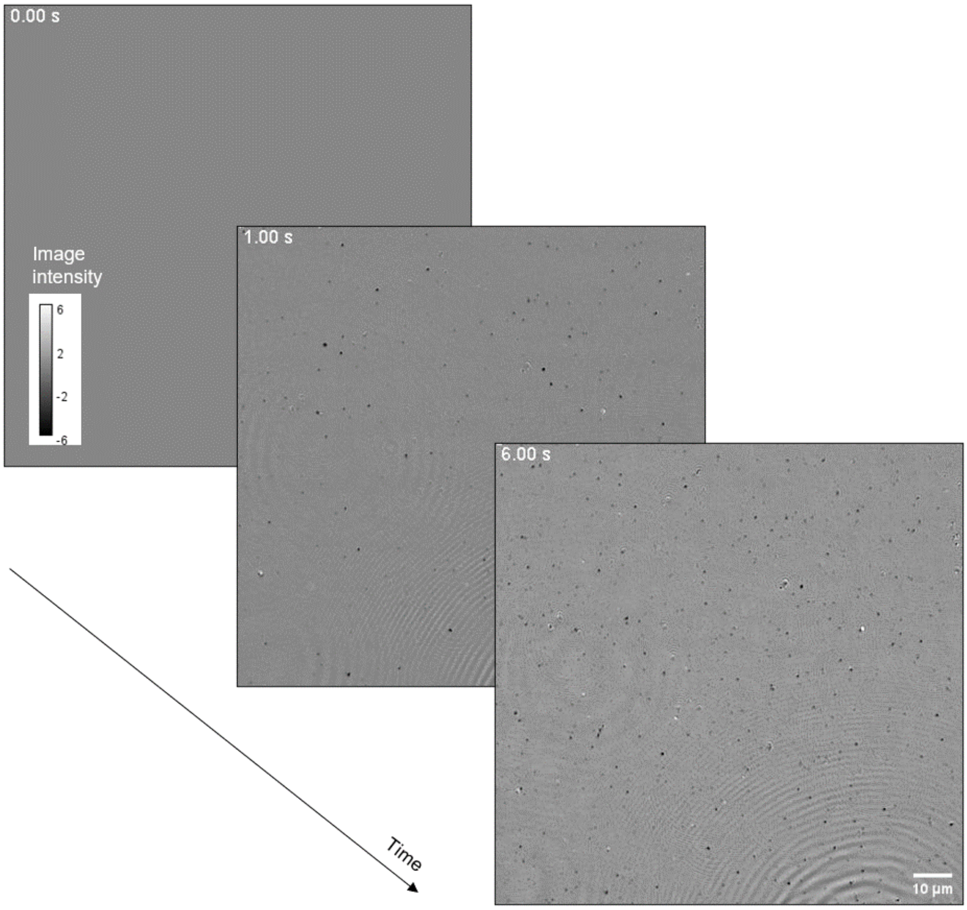


**Supplementary Figure 4. iSCAT images of EVs landing on the surface.** The figure displays snapshots of EVs at three different time points: 0 seconds, 1 second, and 6 seconds.

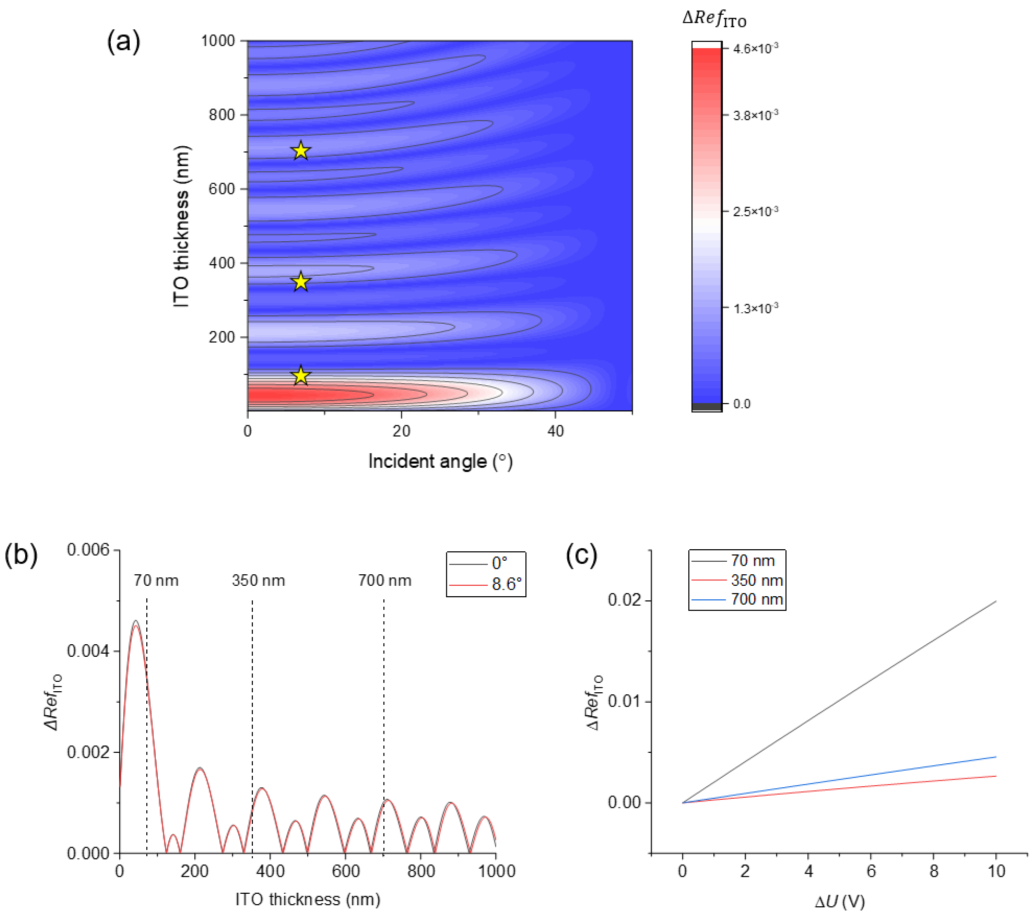


**Supplementary Figure 5. Calculation results of ITO reflectance.** (a) Potential induced reflectance change for ITO with different thickness and different incident light angles, using a 1 V applied potential and assuming no voltage drop across solution resistance. Experimental conditions used in this work are marked by stars in the plot, which are 70 nm, 350 nm, and 700 nm thickness with ~8.6 degrees of incidence angle. (b) Reflectance oscillates with increasing film thickness. The $\Delta Ref_{\mathrm{ITO}}$ values are nearly identical at 0 and 8.6 degrees of incidence, suggesting that slightly tilting the incident angle does not affect iECM sensitivity. $\Delta Ref_{\mathrm{ITO}}$ is highly sensitive to variations in thickness, where minor changes can induce significant reflectance alterations. (c) A plot of $\Delta Ref_{\mathrm{ITO}}$ versus applied voltage demonstrates a linear relationship, aligning with experimental results shown in Figure 2d. However, the magnitude of $\Delta Ref_{\mathrm{ITO}}$ deviates, showing 70 nm as the most sensitive thickness rather than 700 nm. This discrepancy may be due to inaccurate calculation parameters or rapid reflectance fluctuations, but the linear relationship still holds.

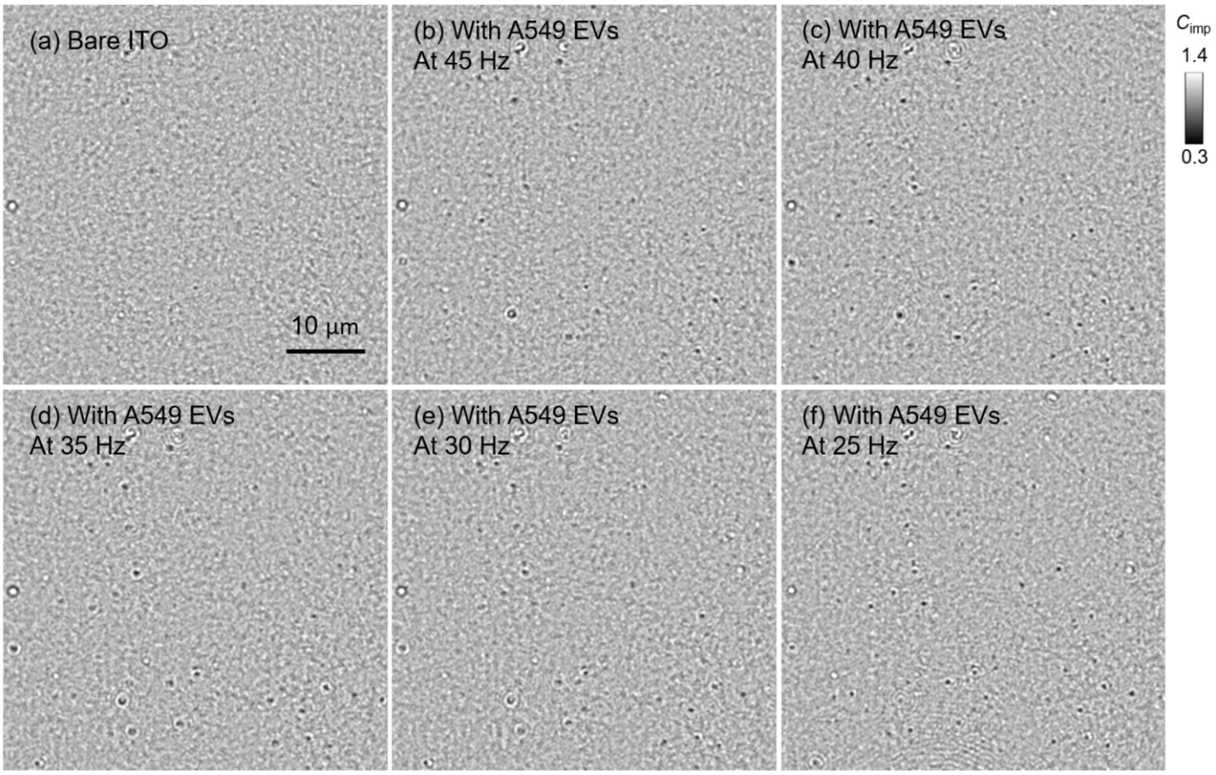


**Supplementary Figure 6. iECM images of an ITO surface before and after adding A549 EVs.** (a) iECM image of the ITO surface before adding EVs. (b-f) iECM images at five different frequencies after adding EVs, where EVs appear as dark spots on the image.

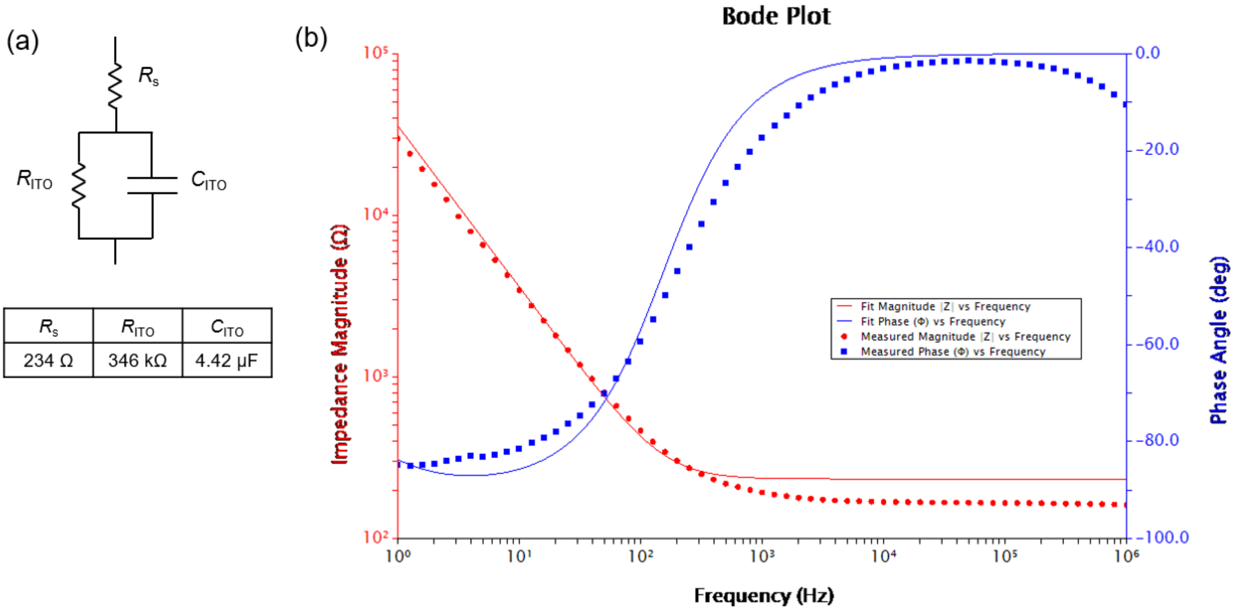


**Supplementary Figure 7. Measuring the impedance of ITO.** (a) The Randle's equivalent circuit used for data fitting, including the measured resistance and capacitance values using traditional electrochemical methods. Note that the exposed ITO area to the solution was 1 cm$^2$. (b) Impedance measurement data collected using a 700 nm ITO coated coverslip and fitted using AfterMath software. The measurements were conducted in 5 times diluted PBS.

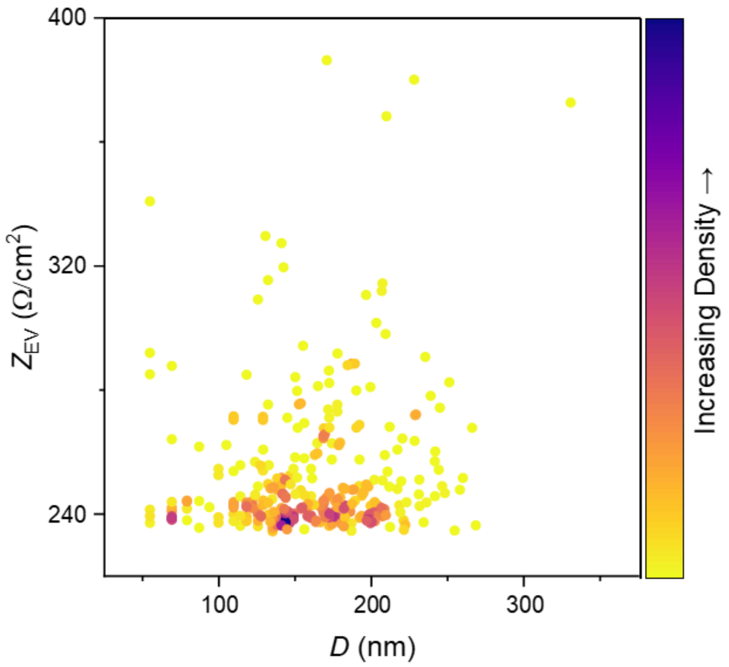


**Supplementary Figure 8. Density map of impedance vs. size for single EVs.** The data is from Figure 3g. The map reveals no obvious clustering.

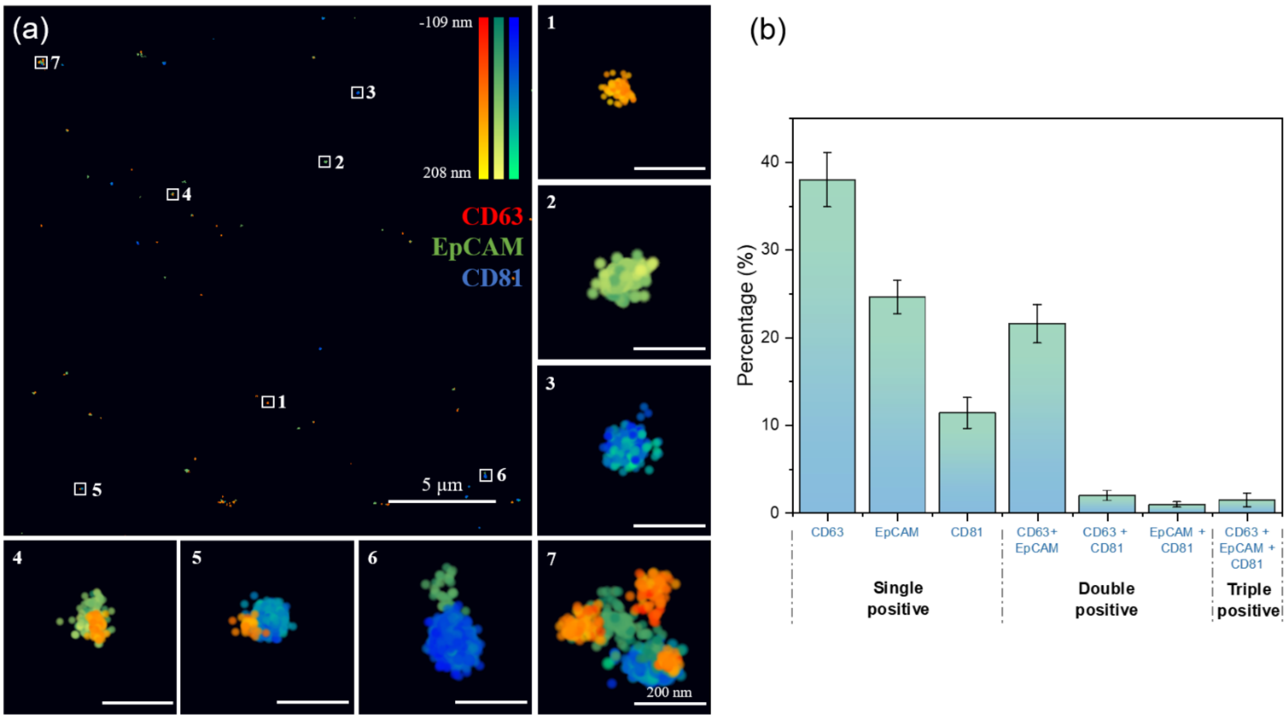


**Supplementary Figure 9. 3D dSTORM imaging of lEVs from A549 cells.** (a) The lEVs were labeled with fluorescent antibodies for CD63, EpCAM, and CD81. Seven representative single EVs (numbered 1-7) are shown enlarged, displaying heterogeneous expression of the proteins, where EVs #1-3 express one marker, #4-6 express two markers, and #7 expresses all three markers. The color bars indicate axial locations. (b) Proportions of lEV subgroups exhibiting single, double, and triple markers. Error bars represent the 95% confidence interval.

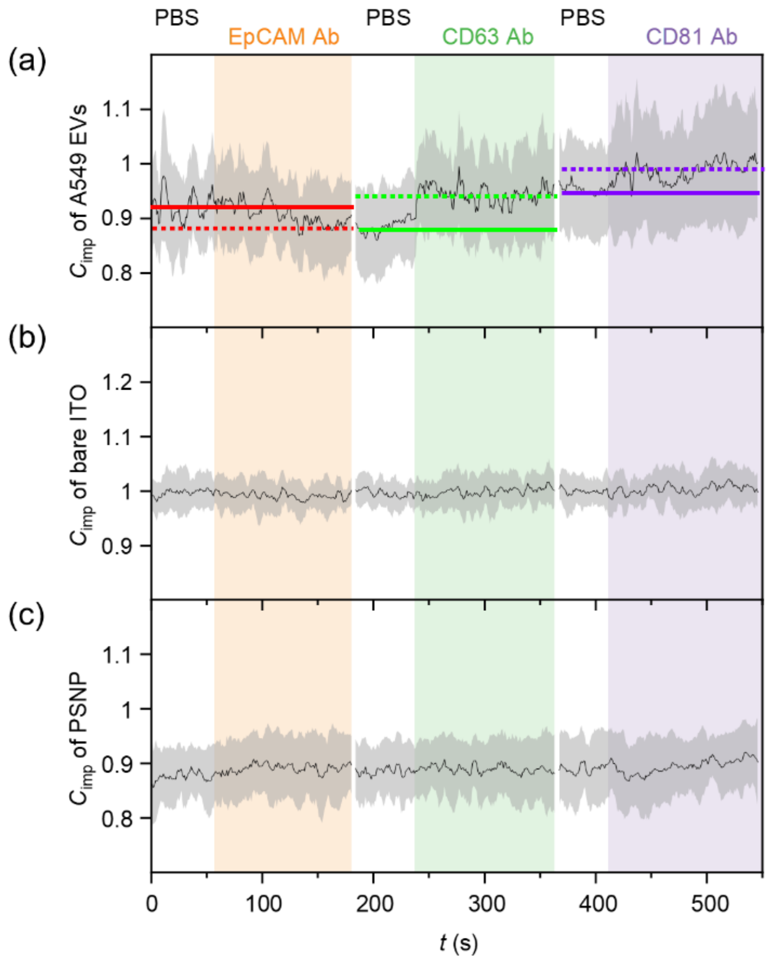


**Supplementary Figure 10. Real-time measurements of antibodies binding to A549 EVs.** (a) Impedance contrast measured from 12 individual EVs was averaged (solid curve), showing a decrease upon EpCAM antibody binding and an increase upon CD63 and CD81 antibody binding. The $C_{\mathrm{imp}}$ levels before and after antibody binding are marked by solid and dashed lines, respectively. The colored areas indicate the incubation periods with the corresponding antibodies. The gray shaded area represents the standard deviation of the 12 EVs. (b) Measurements from 12 bare ITO regions adjacent to the EVs. The curve shows no impedance change and exhibits smaller fluctuations compared to those observed with EVs. (c) Control experiment using PSNPs. The curve shows the average result from 12 single PSNPs, which exhibit no response to the antibodies. The fluctuation level is between that of the EVs and the bare ITO.

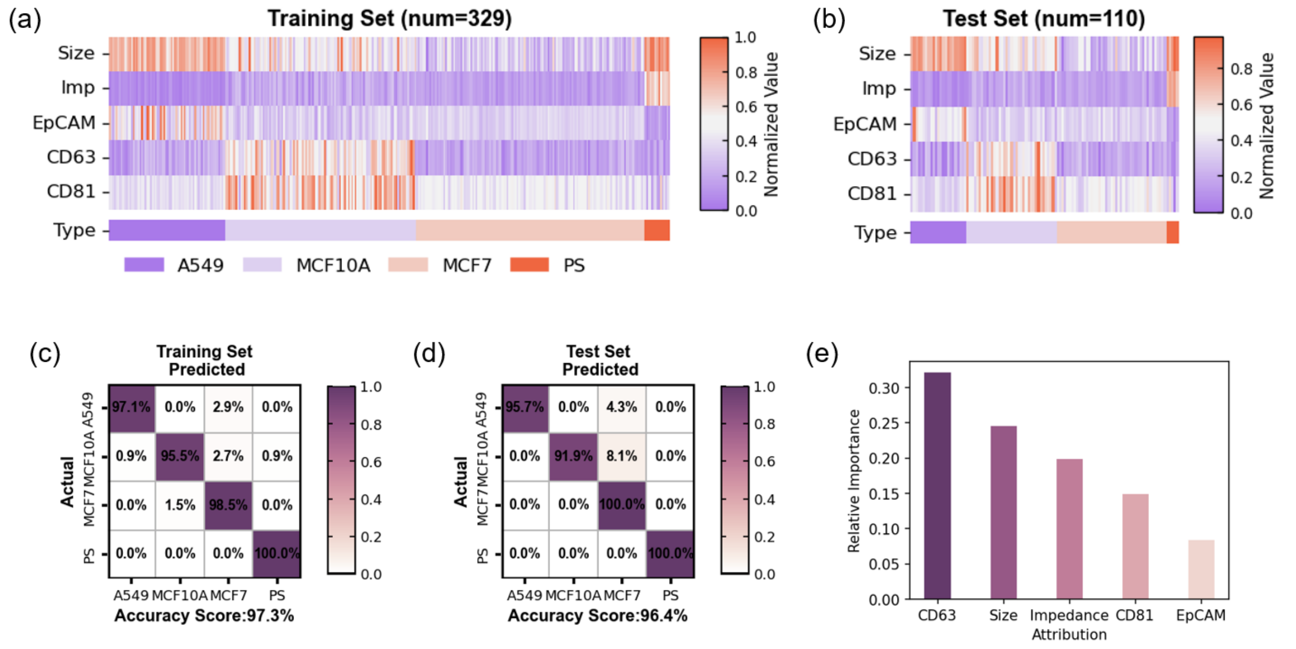


**Supplementary Figure 11. EV classification through Random Forest using 5 metrics (size, impedance at 30 Hz, and 3 protein markers).** (a-b) Heatmap of the training set and test set. (c-d) Confusion matrix of the training set and the test set. (e) Feature importance of Random Forest algorithm. All items add up to 100%. Each data point in the figures represents averaged data from 10 EVs using the bootstrap averaging method.

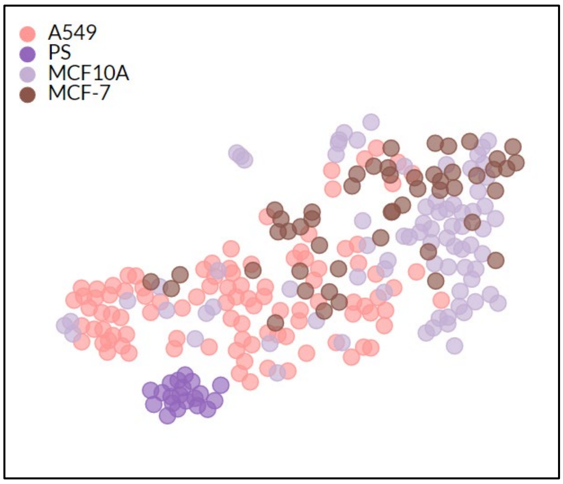


**Supplementary Figure 12. t-SNE plot showing no clear separation between EVs without bootstrap averaging.** Size, impedance at 30 Hz, and antibody binding-induced impedance changes (EpCAM, CD63, and CD81 antibodies) were used as input data.

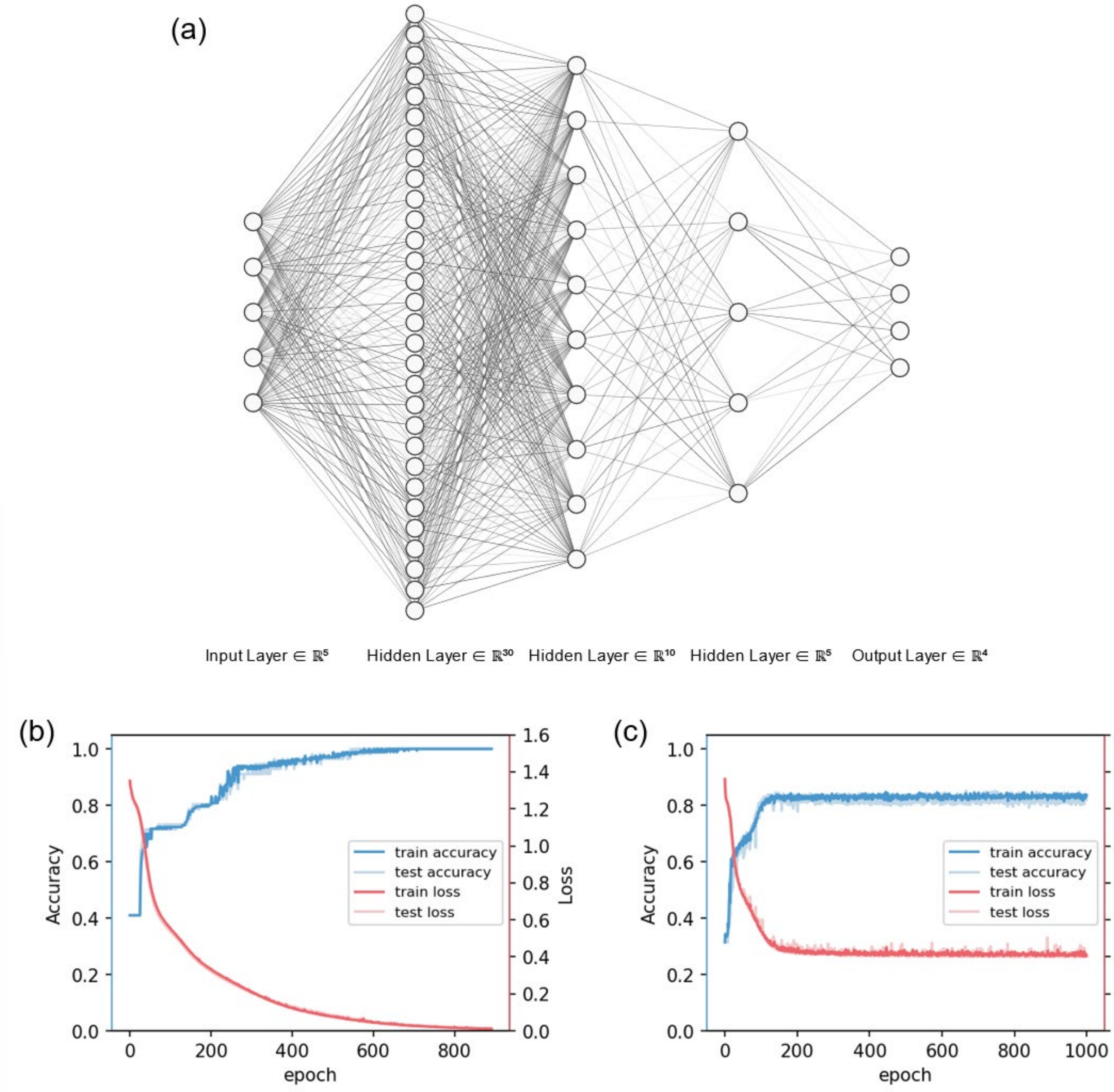


**Supplementary Figure 13. EV classification through Deep Neural Networks.** (a) Schematic diagram of the DNN algorithm used in this work, which has 3 hidden layers with 30, 10 and 5 nodes, respectively. The figure was generated using NN-SVG (https://alexlenail.me/NN-SVG). (b) Accuracy and loss of the training and test sets when using 5 metrics (size, impedance, and 3 protein markers) for classification. The accuracy of both sets reached 100% after 800 epochs. (c) Accuracy and loss of the training and test sets when using impedance fingerprinting spectra (25, 30, 35, 40, and 45 Hz) for classification. The accuracy of both training set and test set reached over 80% after 200 epochs. The input data in (b) and (c) are averaged data from 10 EVs using the bootstrap averaging method.

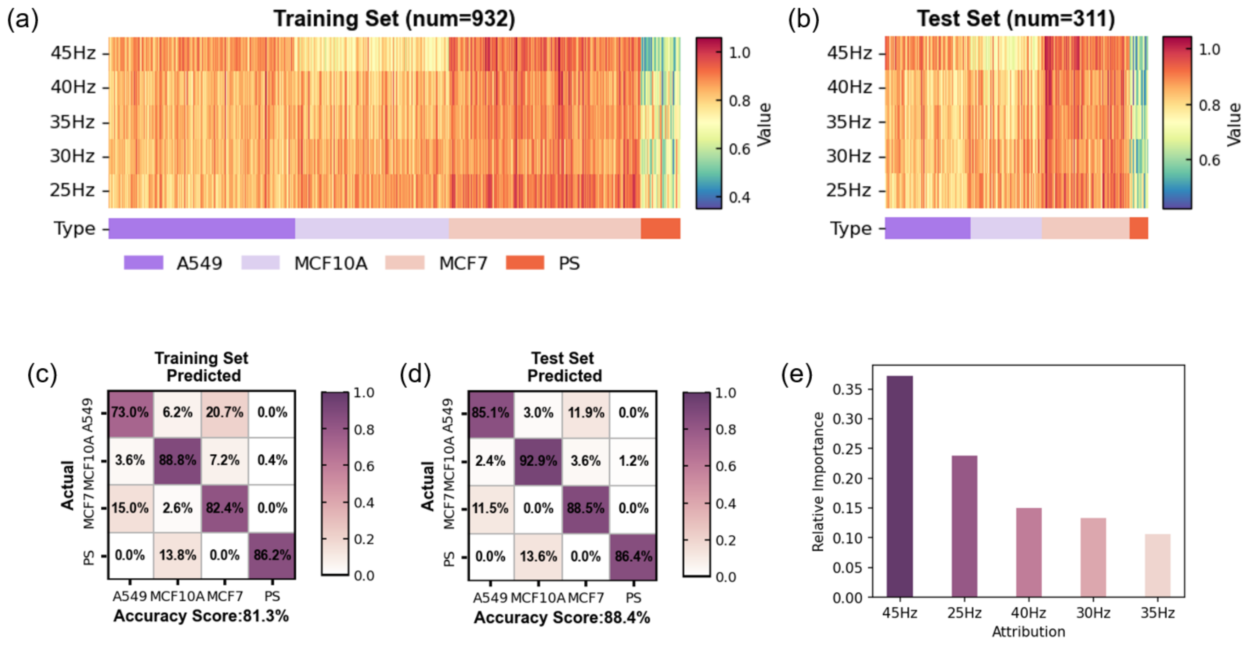


**Supplementary Figure 14. EV classification through Random Forest using impedance fingerprinting spectra (25, 30, 35, 40, and 45 Hz).** (a-b) Heatmap of the training set and test set. (c-d) Confusion matrix of the training set and the test set. (e) Relative importance of Random Forest algorithm. All items add up to 100%. Each data point in the figures represents averaged data from 10 EVs using the bootstrap averaging method.

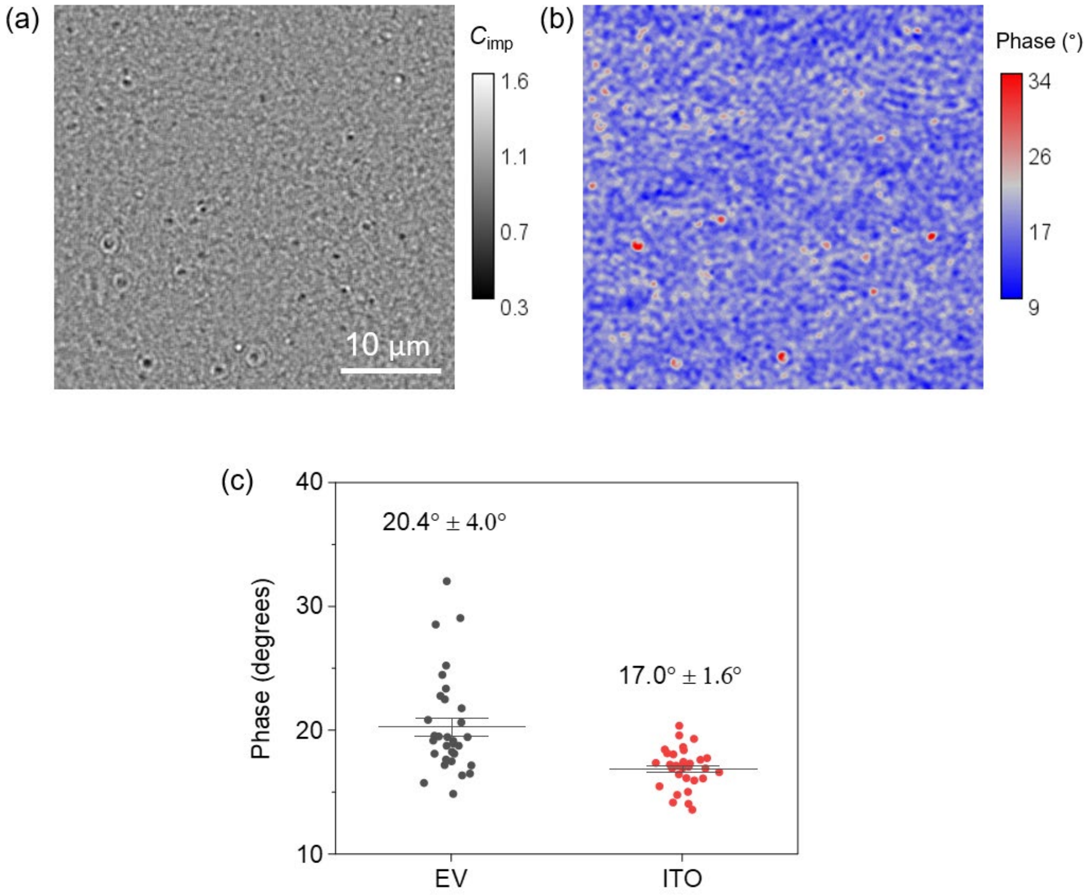


**Supplementary Figure 15. iECM phase images extracted from fast Fourier transform (FFT).** (a) iECM impedance contrast image derived from the FFT amplitude at 30 Hz, showing dark spots as single A549 EVs. (b) The corresponding FFT phase image at 30 Hz. (c) Statistical analysis of the phase of EVs and adjacent background ITO regions. 31 EVs were identified in the amplitude image (a), and the same ROIs were applied to the phase image (b) to measure the phase. 31 background ITO regions in (b) were measured using the same method. The mean values and standard deviations are displayed on the plot, indicating that EVs exhibit only a slightly higher phase (3.4°) than the background, suggesting EVs mainly act as resistors in the equivalent circuit at 30 Hz.

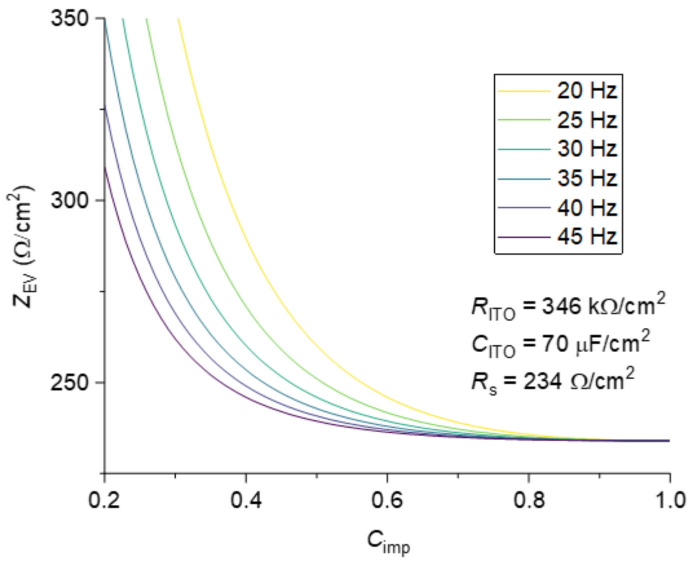


**Supplementary Figure 16. Impedance contrast values of EVs and corresponding impedance.** The impedance values of EVs at different frequencies, calculated using Equation 2 from the main text with known parameter values as indicated in the figure.

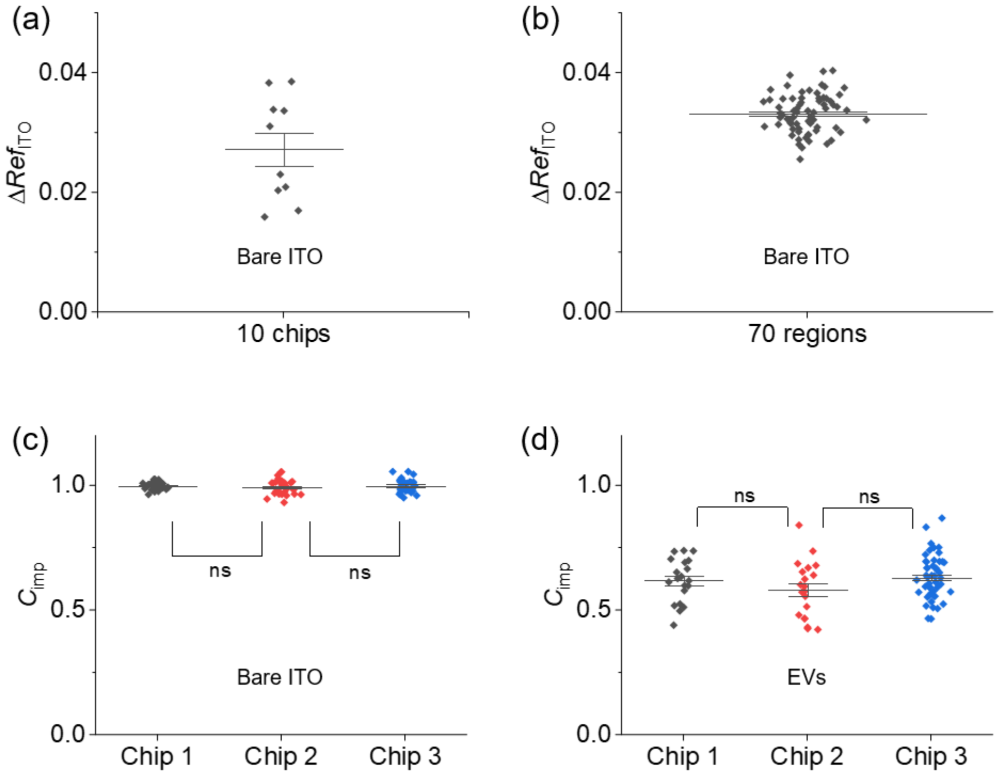


**Supplementary Figure 17. Variations among different ITO chips and regions.** (a) Average $\Delta Ref_{\mathrm{ITO}}$ values obtained from 10 ITO chips of 700 nm thickness. The variation in $\Delta Ref_{\mathrm{ITO}}$ reflects the electrical inhomogeneity within batch of ITO chips. (b) $\Delta Ref_{\mathrm{ITO}}$ values obtained from 70 different diffraction limit-sized regions of interest (ROIs) on the same ITO chip, showing regional electrochemical non-uniformity. (c) Impedance contrast ($C_{\mathrm{imp}}$) measured from three different chips, each analyzed across 30 diffraction limit-sized ROIs. The $C_{\mathrm{imp}}$ variation among different chips are similar. (d) $C_{\mathrm{imp}}$ of A549 lEVs measured from three different chips, showing good reproducibility. $N$ = 20, 20, and 50 for chip 1, 2, and 3, respectively. Errors bars represent the standard deviation in all figures. The applied potential is 10 Vpp at 30 Hz.

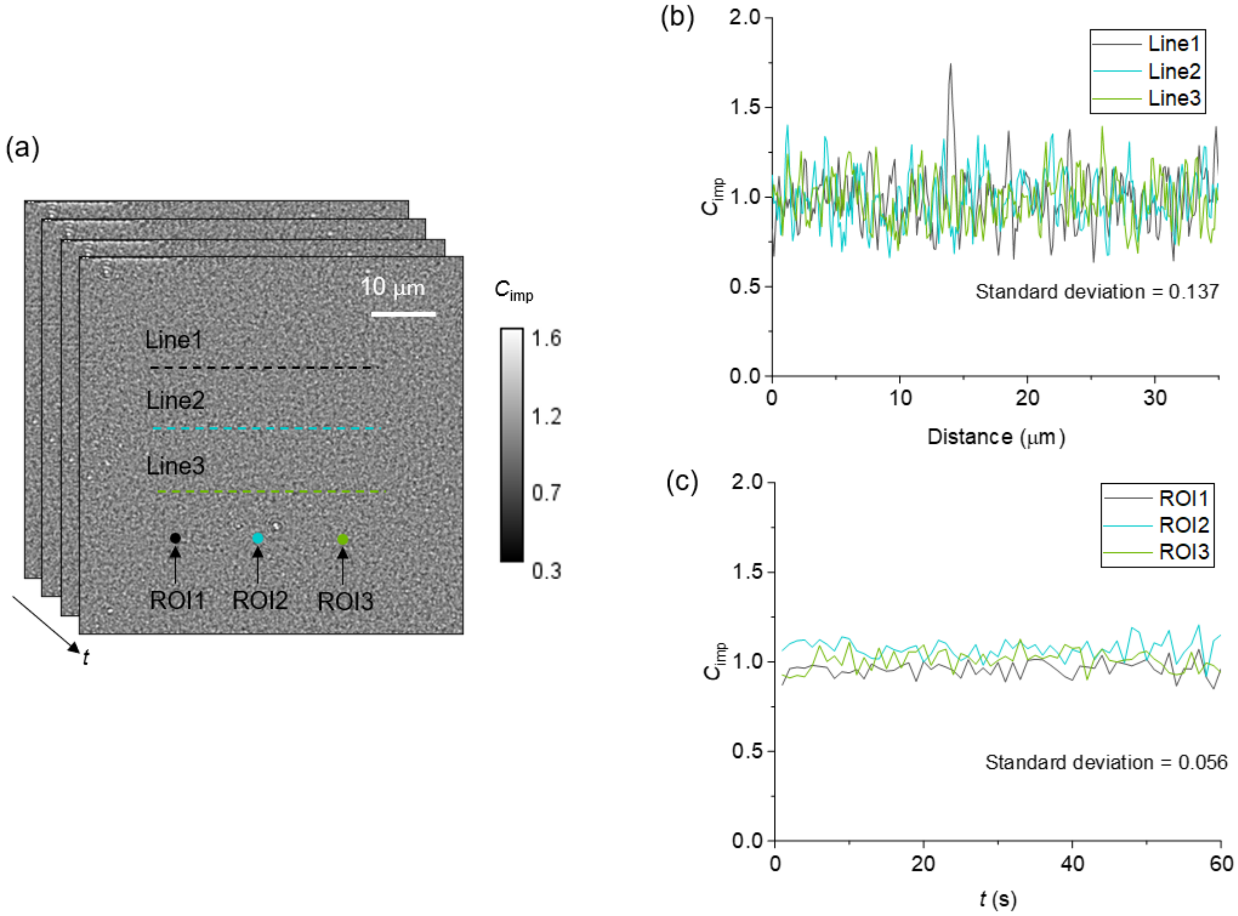


**Supplementary Figure 18. Spatial and temporal noise in impedance imaging.** (a) iECM image sequence with three lines and three diffraction limit-sized ROIs randomly selected for analysis. The applied frequency is 30 Hz. (b) Line profiles showing spatial variations with a standard deviation of 0.137 in terms of $C_{\mathrm{imp}}$. (c) The mean $C_{\mathrm{imp}}$ in the three ROIs are continuously recorded for 60 s. The temporal standard deviation is 0.056, smaller than the spatial noise.

**Supplementary Table 1. Imaging parameters for sizing measurements.**

| Sample | Exposure time (μs) | Frame rate (fps) | Moving average (frames) |
|---|---|---|---|
| 30 nm PSNP | 900 | 1000 | 201 |
| 50 nm PSNP | 1500 | 50 | 51 |
| 75 nm PSNP | 1500 | 50 | 11 |
| 100 nm PSNP | 1500 | 50 | 11 |
| 150 nm PSNP | 1500 | 50 | 11 |
| 200 nm PSNP | 1500 | 10 | 11 |
| lEV and sEV | 1500 | 500 | 51 |

**Supplementary Table 2. Experimental condition for impedance measurements.**

<table>
<tr><th>Experiment</th><th>f (Hz)</th><th>U (V)</th><th>Frame rate (fps)</th><th>Exposure time (μs)</th></tr>
<tr><td rowspan="6">Impedance spectrum measurements</td><td>1</td><td>0.5</td><td>500</td><td>1500</td></tr>
<tr><td>2</td><td>0.5</td><td>500</td><td>1500</td></tr>
<tr><td>5</td><td>1</td><td>500</td><td>1500</td></tr>
<tr><td>10</td><td>2.5</td><td>500</td><td>1500</td></tr>
<tr><td>20</td><td>5</td><td>500</td><td>1500</td></tr>
<tr><td>>20</td><td>10</td><td>500</td><td>1500</td></tr>
<tr><td>Real-time binding detection</td><td>30</td><td>6</td><td>100</td><td>1500</td></tr>
</table>

### Supplementary Note 1. Size conversion factor and detection limit

PSNPs and EVs have different refractive indices, which are 1.59 and 1.35-1.55 (we use 1.42 in this work), respectively. Therefore, the size of EVs determined using the PSNP calibration curve requires adjustment for the refractive index difference. For Rayleigh scattering, the polarizability $\alpha$ is defined as,

$$\alpha = \frac{1}{2}\pi D^3 \left(\frac{{n_\mathrm{p}}^2 - {n_\mathrm{m}}^2}{{n_\mathrm{p}}^2 + 2{n_\mathrm{m}}^2}\right) \tag{S1}$$

Where $D$ is the particle diameter, $n_\mathrm{p}$ is the refractive index of the nanoparticle (or EV), and $n_\mathrm{m}$ is the refractive index of the medium (PBS, $n_\mathrm{m} = 1.334$). For a PSNP and an EV of the same size, $\alpha_\mathrm{PS} = 2.90\alpha_\mathrm{EV}$, and consequently, $D_\mathrm{EV} = 0.70D_\mathrm{PS}$.

The signal-to-noise ratio (SNR) for a 50 nm PSNP is approximately 12, as shown in Figure S2a. Considering an SNR threshold of 3 for reliable signal detection above background noise, the smallest detectable PSNP size is calculated to be 32 nm, which corresponds to an EV size of 45 nm. A notably low SNR is observed for the 30 nm PSNPs sample (close to the detection limit), which has an actual measured diameter of 33.7 nm by dynamic light scattering (DLS).

### Supplementary Note 2. Theory of impedance imaging

We model the ITO film as a conductive layer with a frequency-dependent dielectric constant $\varepsilon_{ITO}(\omega)$, described by the Drude model,

$$\varepsilon_{ITO}(\omega) = 1 - \frac{n_e e^2}{\varepsilon_0 m_e \omega^2} \tag{S2}$$

where $n_e = 5 \times 10^{20}\ \text{cm}^{-3}$ is the electron density,[2] $e = 1.6 \times 10^{-19}\text{C}$ is the electron charge, $m_e$ is the electron mass, $\varepsilon_0$ is the vacuum permittivity, and $\omega$ is the angular frequency. The surface charge density $\sigma$ of the film (thickness $d_{ITO}$) can be moculated by applying potential $(U)$, and its change $\Delta\sigma$ is given by

$$\Delta\sigma = c\Delta U = -ed_{ITO}\Delta n_e \tag{S3}$$

where $c$ is the capacitance density of ITO and $\Delta U$ is the potential change. Combining Equations S2 and S3, we have:

$$\Delta\sigma = c\Delta U = -\frac{ed_{ITO}n_e}{\varepsilon_{ITO} - 1}\Delta\varepsilon_{ITO} \tag{S4}$$

and

$$\Delta\varepsilon_{ITO} = \frac{c(1 - \varepsilon_{\text{ITO}})}{ed_{ITO}n_e}\Delta U \tag{S5}$$

where $\frac{c(1-\varepsilon_{ITO})}{ed_{ITO}n_e} = \beta$ (which corresponds Equation 2 in the main text). The capacitance density of ITO in 5 times diluted PBS is measured to be $c \sim 5.0\ \mu\text{F/cm}^2$, and $\varepsilon_{\text{ITO}} = 4$ according to the manufacturer's datasheet. Thus, the calculated values of $\beta$ for $d_{ITO}$ = 70, 350, and 700 nm are -0.0268, -0.0054, and -0.0027 $\text{V}^{-1}$, respectively. The refractive index of ITO as a function of potential change is expressed as:

$$n_{\text{ITO}}(\Delta U) = \sqrt{\varepsilon_{\text{ITO}}(\Delta U)} = \sqrt{4 + \beta\Delta U} \tag{S6}$$

The reflectivity of the ITO coated glass coverslip can then be simulated using Winspall software and plotted against $\Delta U$. The result is shown in Figure S5c.

### Supplementary Note 3. EVs exhibit resistive behavior

In this study, the impedance of EVs is modeled as a combination of capacitance and resistance, where the significance of each component in the equivalent circuit depends on the applied frequency. Below 1 MHz, the capacitance of the lipid bilayer is substantial, making the EVs effectively insulating.[3] Consequently, the EV primarily behaves as a resistor. Using FFT analysis (Methods), we extracted phase information from the iECM images and observed that the presence of EVs introduces only a 3.4° phase change (Figure S15). This confirms the predominantly resistive behavior of EVs at low frequencies. As the frequency increases, the membrane begins to exhibit more capacitive behaviors, and the total impedance becomes a combination of both capacitance and resistance components. At very high frequencies, most of the current flows through the capacitive component of the EV. Under these conditions, iECM becomes particularly effective for measuring membrane changes, making it an excellent tool for membrane protein profiling. However, impedance imaging at high frequencies is challenging, because most commercial CMOS cameras cannot achieve the necessary high frame rates. Additionally, applying high frequencies requires the miniaturization of electrodes to ensure rapid response times. For instance, a modulation frequency of 10 MHz would require microelectrodes as small as 40 μm.[3] Such requirements demand meticulous design of the sensor chip, including precise electrode patterning and sophisticated sample handling strategies.

### Supplementary Note 4. Simulation parameters

Parameters used in Figure 3d include $R_{\mathrm{ITO}} = 346\ \mathrm{k\Omega}$, $C_{\mathrm{ITO}} = 70\ \mathrm{\mu F}$, $R_{\mathrm{s}} = 234\ \Omega$, $R_{\mathrm{EV}} = 50 - 200\ \Omega$, and $C_{\mathrm{EV}} < 100\ \mathrm{\mu F}$. Changing the values of $R_{\mathrm{EV}}$ and $C_{\mathrm{EV}}$ within the indicated range will not influence the shape of the curves significantly. The values for these parameters are determined as follows: $R_{\text{ITO}}$ and $R_{\text{s}}$ are measured values obtained through traditional electrochemistry and are treated as known constants. Then $C_{\text{ITO}}$ is determined to be 70 μF by fitting the "ITO" data in the figure. This value is higher than the 4.42 μF measured by conventional electrochemistry methods (Figure S7), due to the different measurement methods (optical vs. electrical). Using $R_{\text{ITO}}$, $C_{\text{ITO}}$, and $R_{\text{s}}$ as inputs, we fitted the "ITO+EV" data in Figure 3d. Due to the narrow frequency window and the level of variation in $\Delta Ref$, accurate values for $R_{\text{EV}}$ and $C_{\text{EV}}$ could not be precisely extracted, only a range could be estimated with $R_{\text{EV}} = 50 - 200\ \Omega$ and $C_{\text{EV}} < 100$ μF. This means we can only obtain the impedance value from the measurement, not the precise $R_{\text{EV}}$ and $C_{\text{EV}}$. The $C_{\text{imp}}$ curve in Figure 3d, which shows the ratio between "ITO+EV" and "ITO", confirmed alignment with experimental observations. Thus, we adopted $R_{\text{ITO}} = 346\ \text{k}\Omega$, $C_{\text{ITO}} = 70\ \mu\text{F}$, and $R_{\text{s}} = 234\ \Omega$ for calculating the impedance of single EVs in Figure 3g.

**Supplementary Note 5. EV characterization**

*Transmission Electron Microscopy (TEM).* 5 μL EV sample was dropped onto a TEM grid, negatively stained with 5 μL 1% sodium phosphotungstate, and allowed to dry at room temperature for 15 minutes. Then the EVs were imaged using a Jeol JEM-1400flash (JEOL Ltd., Japan).

*Nanoparticle Tracking Analysis (NTA).* EVs were diluted to the appropriate concentration in 1×PBS and analyzed using a ZetaView instrument (Particle Metrix, Germany) for size distribution and concentration. Data processing was done using ZetaView software (version 8.05.14 SP7).

*Western Blot.* EV proteins were denatured by boiling and separated using electrophoresis (Simple PAGE 4-12%

Bis-Tris SDS-PAGE Gel, Sangon Biotech., China). Proteins were transferred to a PVDF membrane (Millipore, USA) at 300 mA for 45 minutes. The membrane was blocked with 5% skim milk and washed thrice with 1×TBST buffer (25 mM Tris, 0.15 M NaCl, 0.05% Tween-20, pH 7.5) for 10 minutes each. Overnight incubation at 4°C was done with primary antibodies (Proteintech, China) for CD63 (1:500), TSG101 (1:4000), CD81 (1:2000), Alix (1:3000), Calnexin (1:20000), EpCAM (1:1000), and GAPDH (1:10000). After rinsing three times with 1×TBST, the PVDF membrane was incubated with corresponding secondary antibodies, HRP-conjugated Affinipure Goat Anti-Rabbit (Cell Signaling Technology, MA, USA) / Goat Anti-Mouse (Proteintech, Wuhan, China) IgG(H+L) at room temperature for 1 h. Finally, membranes were washed, exposed to ECL reagents (Biosharp, China) for 1 minute, and imaged using a JS-M6P chemiluminescence imaging system (Peiqing Tech., China).

*Direct Stochastic Optical Reconstruction Microscopy (dSTORM).* The coverslips were cleaned with ethanol and DI water for three times. After a 10-minute incubation for EV adsorption, the coverslips were flushed three times with 1×PBS to remove unbound EVs and blocked with 1% BSA in PBS for 30 minutes. EVs were then incubated with an antibody mixture containing Alexa Fluor 647 labeled CD63 antibody (1:80 in 1% BSA, 2.5 μg/mL), Alexa Fluor 488 labeled CD81 antibody (1:400 in 1% BSA, 0.125 μg/mL), and Alexa Fluor 555 labeled EpCAM antibody (1:2000 in 1% BSA, 0.015 μg/mL) at room temperature for 1 hour. After washing away excess antibodies with 1×PBS, 60 μL dSTORM imaging buffer (a mixture of glucose buffer, glucoamylase and β-mercaptoethanol (BME) at a ratio of 100:1:1, dSTORM smart kit, Abbelight) was added, and the coverslip, with antibody-labeled EVs, was sealed on a slide for imaging. Imaging was performed on a single-molecule localization microscope (SAFe 180, Abbelight) under TIRF mode, followed by reconstruction and data processing using NEO Analysis software.
